# Voltage Imaging of CA1 Pyramidal Cells and SST+ Interneurons Reveals Stability and Plasticity Mechanisms of Spatial Firing

**DOI:** 10.1101/2025.08.20.671230

**Authors:** Rotem Kipper, Yaniv Melamed, Qixin Yang, Gal Shturm, Shulamit Baror-Sebban, Yoav Adam

## Abstract

Hippocampal place cells (PCs) are important for spatial coding and episodic memory. PCs’ representations are modulated upon transitioning between environments (global remapping) but also change with repeated exposure to familiar spaces (representational drift). To gain insights into the mechanistic basis for this unique balance between circuit plasticity and stability, we used voltage imaging to longitudinally record the subthreshold and spiking activity of pyramidal neurons (PNs) and somatostatin-positive (SST) interneurons in CA1 during virtual navigation. A fraction of cells from both populations showed spatial representations, but many SSTs were speed-tuned or fired uniformly across space. Intracellular recordings revealed increased theta power and asymmetric ramp-like depolarization in PN place fields, while SSTs exhibited symmetric depolarization with no theta increase. Longitudinal recordings across weeks demonstrated representational drifts in both populations, although SSTs exhibited remarkably stable firing and subthreshold properties. Transition to a novel environment induced remapping in both populations, accompanied by increase in SST activity and reduction in PNs. These results provide new insights into how hippocampal circuits balance representational stability with experience-dependent plasticity.

## Introduction

Hippocampal place cells are excitatory PNs that encode an allocentric map of space by exhibiting elevated firing rates at specific locations (’place fields’)^1^. This spatial coding is considered an essential substrate for the formation of spatial and episodic memories^2–4^. Two key properties of this code pertain to its dynamic nature across environments and over time. In any given environment, only a subset of CA1 PNs are active place cells, resulting in a sparse population code^5^. Upon transitioning to a different environment, this active ensemble undergoes a reorganization, termed global remapping, where place fields reconfigure largely randomly^6–8^. Even within a single environment, hippocampal representations are not static; instead, they exhibit gradual changes, or representational drift, driven by the passage of time and ongoing experience^9–11^.

Recent intracellular studies have identified a plasticity mechanism operating on behavioral timescales, termed behavioral time-scale synaptic plasticity (BTSP)^12–14^. BTSP enables the rapid formation of place fields following the occurrence of a dendritic plateau potential triggered by coincidental input from the entorhinal cortex (EC) and CA3. Depolarization by the plateau spreads throughout the neuron and potentiates spatially selective inputs from CA3 that were active around the time of the plateau, thereby forming a place field at its position^15^. The responsible input pathways from CA3 and EC are both dynamically modulated by a diverse population of inhibitory interneurons, which target distinct dendritic compartments of PNs^16,17^ with precisely timed activation patterns^18,19^. This circuit architecture enables fine-tuned, compartment-specific control over synaptic inputs and plasticity, suggesting that the formation of place fields, as well as the dynamics of their remapping and drift, are critically dependent on the local microcircuit in which CA1 PNs are embedded.

Somatostatin-expressing (SST+) neurons are of particular interest regarding remapping and place field formation. They include oriens-lacunosum moleculare (OLM) cells that innervate the distal apical tuft and regulate integration of EC input^20,21^, potentially influencing BTSP induction^22^, and bistratified cells that innervate the basal dendrites and modulate inputs from CA3^21,23^. Several recent calcium imaging studies have reported a decrease in SST activity in novel environments, suggesting that transient disinhibition of PN dendrites facilitates BTSP and promotes the formation of new place fields^24–28^. However, calcium imaging provides a relatively slow proxy for neuronal activity, therefore lacking precise spike timing resolution, and is limited in reporting activity of high-firing neurons^29,30^ such as SSTs^31,32^. Furthermore, while we have some understanding of the plasticity rules driving place cell formation in PNs based on intracellular studies^12,13^, the plasticity mechanisms underlying feature selectivity in interneurons are mostly elusive. Voltage recordings at high temporal resolution are therefore required for resolving the dynamics of inhibitory interneurons during global remapping.

Studies on CA1 representational drift have focused almost exclusively on PNs using either extracellular recordings or calcium imaging^9,33–36^, and none have specifically targeted distinct interneuron subtypes. Consequently, the contribution of different interneuron classes to representational drift in PNs, and the extent to which interneurons themselves exhibit drift, remains largely unexplored. Furthermore, data on the long-term stability of spiking and subthreshold activity in PNs and interneurons is still lacking. This gap arises from the complementary limitations of existing techniques: calcium imaging lacks access to subthreshold membrane dynamics and is poorly suited for capturing activity in high-firing interneurons, while intracellular electrophysiology offers high temporal resolution but is low-throughput, difficult to apply to molecularly defined subtypes^19,37^, and not compatible with longitudinal recordings.

Genetically encoded voltage indicators (GEVIs) offer a complementary approach that combines the precision of intracellular electrophysiology with the scalability, cell-type specificity, and longitudinal capabilities of optical imaging^31,38^. Here, we performed voltage imaging with a near-infrared GEVI in head-fixed mice navigating a one-dimensional (1D) virtual environment to obtain a large dataset of intracellular voltage recordings from CA1 pyramidal cells and SST+ interneurons across weeks and during context switching. This approach enabled direct comparison of the spiking and subthreshold determinants of spatially tuned cells across both cell types and uncovered differences in the subthreshold correlates of their spatial tuning. In addition, we found that SST cells can be classified into at least 3 tuning types – place cells, speed cells, and spatially uniform cells. By tracking the same cells across weeks, we found that representations in both PNs and SST+ interneurons drifted over time, even though SST+ cells exhibited remarkably stable firing and subthreshold properties. Finally, exposure to a novel environment triggered remapping of spatially tuned cells from both populations, accompanied by a transient increase in SST+ cell firing and a corresponding decrease in PN activity. Together, these findings shed light on the role of SST+ interneurons in both stability and plasticity mechanisms within the CA1 microcircuit and highlight voltage imaging as a powerful tool for longitudinal, cell-type-specific recordings with intracellular resolution at scale.

## Results

### Cell-type specific voltage imaging during virtual reality navigation

To investigate the supra- and subthreshold dynamics of distinct CA1 cell types during behavior, we performed voltage imaging through an imaging cannula in head-fixed mice running in a one-dimensional (1D) virtual reality (VR) environment^39^ for 10% sucrose rewards. Rewards were delivered at a fixed reward zone with distinct spatial features. VR training and imaging were performed in a custom-designed compact VR setup that can be easily integrated under standard upright microscopes (Fig. 1a, Methods). We expressed the near-infrared GEVI somArchon1^38^ in either pyramidal neurons (PNs) or SST+ interneurons (SST INs), by injecting Cre-dependent AAV vectors into mice expressing Cre-recombinase in CaMKII+ or SST+ cells, respectively. To achieve the sparse expression in PNs necessary for 1-photon voltage imaging in the densely packed pyramidal cell layer, we employed an intersectional strategy by injecting CaMKII-Cre mice with a mixture of Flp-dependent Archon with low titer Cre-dependent Flp virus to restrict expression to doubly infected PNs^32^ (Fig. 1a).

**Fig. 1:**
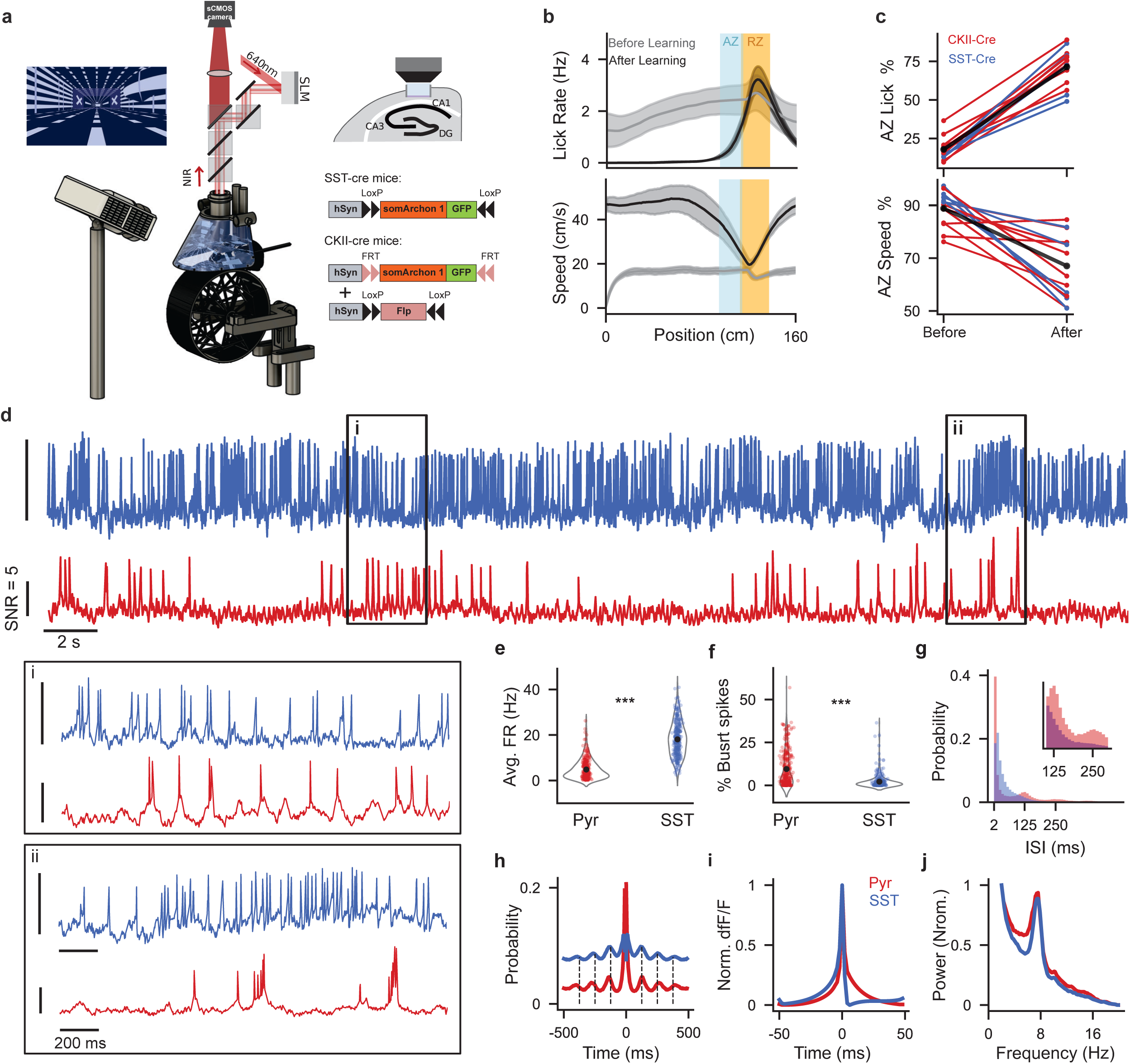
Spiking and subthreshold activity in Pyramidal cells and SST+ interneurons. **a**, Left - optical system for 1-photon voltage imaging using targeted illumination, combined with a custom VR system for head fixed mice. Right - Illustrations of the cannula under the objective, and viral strategy for expressing somArchon1 in SST or CKII-cre mice. **b**, Top: Lick rate by position before and after learning the task for a single mouse. Bottom: Same for running speed. AZ - anticipation zone, RZ - reward zone. **c**, Top: Percentage of licks within the AZ for all mice, CKII-Cre in red, SST-Cre in blue. Bottom: avg. speed within AZ as a percentage of maximal average speed (N = 9 CKII-Cre and 5 SST-Cre mice). **d**, Optically recorded voltage traces (20 seconds) from a representative CA1 SST cell (blue) and PN (red). Vertical scale bars indicate 5 STDs of the fluorescence signals. Insets – zoom in on 2 seconds at two indicated locations. **e**, Average firing rates of all recorded cells (mean ± SEM = PNs: 4.81 ± 0.25, SSTs: 18.07 ± 0.43; P = 3.00 X 10^-80^, Wilcoxon rank-sum test. N = 232 PNs and 187 SSTs). **f**, Percentage of spikes that have a neighboring spike within ≤ 10ms (mean ± SEM = PNs: 9.52 ± 0.6, SST INs: 2.17 ± 0.24, P = 7.37 X 10^-26^, Wilcoxon rank-sum test). **g**, Distribution of inter-spike-intervals (ISIs) across all cells of each population (inset; zoomed in to ISIs >100ms). **h**, Population average auto correlograms. Dashed lines highlight the peaks at 125 ms multiplicands away from the spike, corresponding to theta-modulated firing at around 8 Hz. **i**, Normalized average spike shapes computed on all spikes (excluding spikes with another spike within ± 50 ms) from each population (N=75,604 spikes for CKII-cre and 58,775 for SST-cre mice) **j**, Normalized population average power spectral density of the subthreshold Vm trace displays a prominent peak at ∼ 7.3 Hz in both populations.

To achieve voltage imaging at a high signal-to-noise ratio (SNR), we implemented our previously published targeted illumination approach^31^, minimizing background fluorescence, photobleaching, and sample heating. To this end, we used spatial light modulator (SLM)-based holographic illumination of selected cells^32^ (Fig. 1a, Methods). Mice underwent 2–3 weeks of daily training on the running wheel (see Methods) within a single virtual environment until they were able to complete 10 laps per minute. All mice from both strains learned the task as indicated by increased licking and reduced speed in the reward anticipation zone (Fig. 1b,c). Once this criterion was met, the first imaging session was conducted in the same familiar environment, followed by at least one additional imaging session in the same familiar environment. In their last imaging session, each trial began in the familiar space and transitioned midway to a novel, unfamiliar one. Using this paradigm, we first compared neuronal activity during spatial navigation across the two populations. Next, we analyzed their stability upon repeated exposure to the same environment across weeks and their responses to a context switch.

In the first session, we performed intracellular voltage recordings from 118 PN FOVs with a total of 232 active PNs and 77 SST FOVs with 187 active SST cells (Extended Data Fig. 1) in 9 CaMKII-Cre mice and 5 SST-Cre mice. For statistical analyses, all recorded cells from each population were pooled together. Cells from both populations exhibited distinct spiking and subthreshold dynamics. SST cells displayed distinctly high firing rates while PNs fired more sparsely, displaying a bursty, theta-modulated firing pattern (Fig. 1e-h) and distinct spike waveforms (Fig. 1i). In addition, the power spectra of the subthreshold activity in both populations displayed prominent intracellular theta oscillations, peaking at ∼7.3 Hz (Fig. 1j).

Analyzing their activity across space (Fig. 2a), we found that in agreement with previous studies, approximately 30% of PNs^9,40,41^ and 20% of SST+ interneurons^24^ showed increased firing rates in specific positions along the track, classifying them as spatially tuned (Fig. 2b and Extended Data Fig. 2, see Methods). Notably, cells from both populations tiled the entire track but exhibited opposing spatial biases: PNs overrepresented the reward zone, whereas SST+ INs underrepresented it (Fig. 2c,d and Extended Data Fig. 2c). Consistent with previous results^42^, place fields of both populations were comparable in spatial information, width, and reliability (Fig. 2f-g). However, they differed in their in-field to out-of-field firing rate ratios (Fig. 2i), reflecting the significantly lower baseline firing rates of PNs outside of their place fields.

**Fig. 2:**
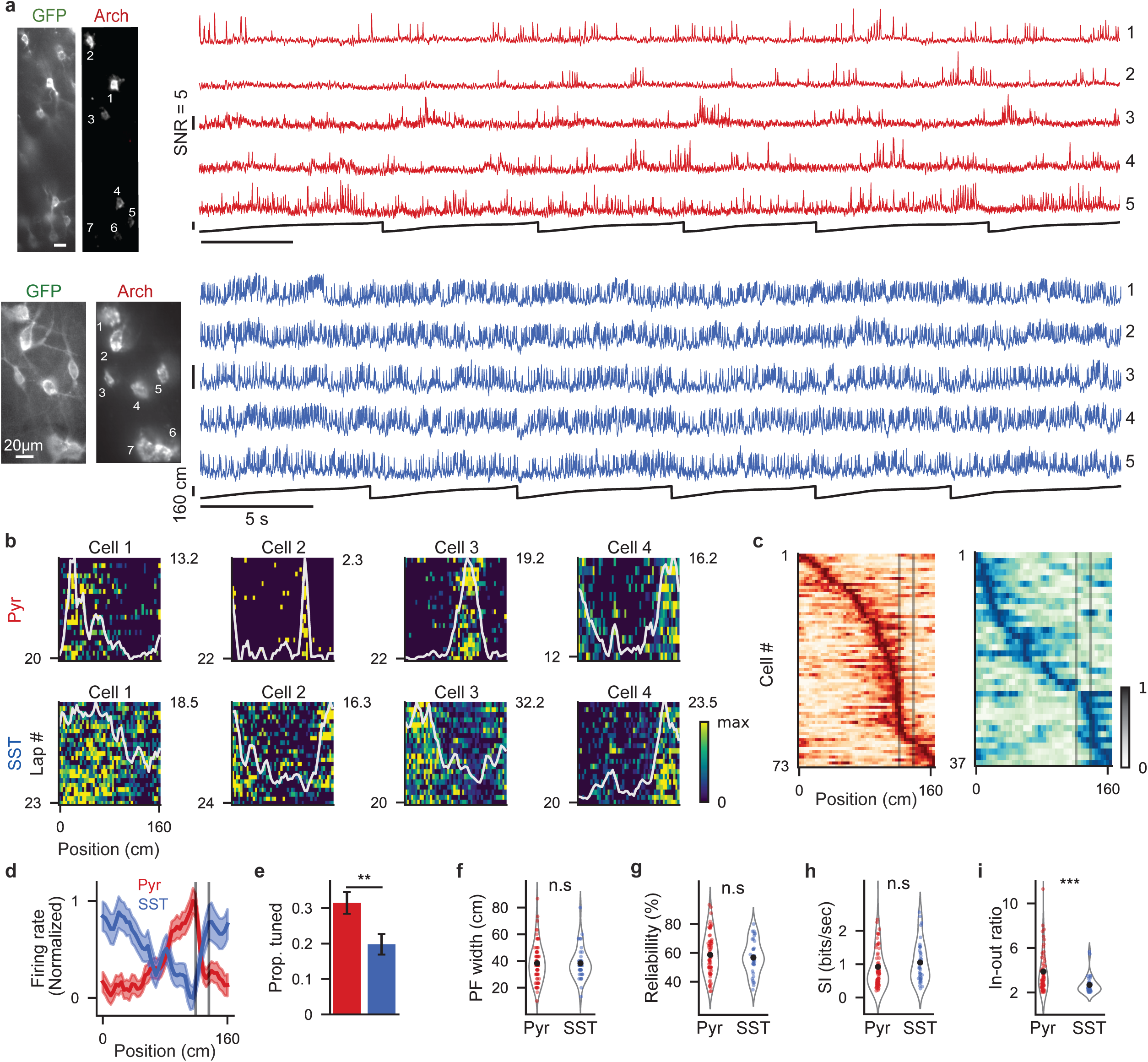
Spatial tuning of SSTs and PNs during VR spatial navigation. **a**, Two fields of view recorded in CKII-Cre (top) or SST-Cre (bottom) mice under wide-field blue LED illumination (GFP) or 639nm SLM-patterned holographic illumination (Arch). For each FOV, voltage traces of 5 simultaneously recorded cells are presented. Black line - mouse virtual position. **b**, Firing rate (FR) by position heatmaps of 4 spatially tuned cells from each population. Gray line denotes the average FR across all laps. Top right - maximal value of average FR vector (Hz), bottom left - total number of laps in the trial. **c**, Average FR by position vectors of all spatially tuned cells from both populations, sorted by the location of their peak. Each vector (row) is normalized from 0 to 1. Gray lines denote the reward zone. **d**, Population averages for all cells in c. Shaded areas indicate ± SEM. **e**, Proportion of spatially tuned cells out of total recorded (Error bars indicate ± SEM. PNs: 73/232, SSTs: 37/187, P = 0.007, Two proportions Z-test). **f**, Place field widths in cm (mean ± SEM = PNs: 38.31 ± 1.66, SSTs: 38.29 ± 2.03, P = 0.95, Wilcoxon rank-sum test). **g**, Percentage of laps with significant firing within the place field (mean ± SEM = PNs: 58.6 ± 1.67, SSTs: 56.8 ± 1.94, P = 0.49, independent t-test). **h**, Spatial information rate of the average firing rate vector in bits/second (mean ± SEM = PNs: 0.92 ± 0.09, SSTs: 1.05 ± 0.1, P = 0.07, Wilcoxon rank-sum test). **i**, In-field to out-field firing rate ratio. (mean ± SEM = PNs: 3.89 ± 0.27, SSTs: 2.68 ± 0.14, P = 2.9 X 10^-5^, Wilcoxon rank-sum test).

In summary, by performing voltage imaging in both cell populations under the same behavioral paradigm, we obtained the first large-scale intracellular dataset from SST+ interneurons, enabling a direct comparison of their firing and spatial tuning properties to those of PNs. To further characterize these differences, we next analyzed the subthreshold dynamics underlying their spatial firing.

### Distinct subthreshold signatures of SST and Pyramidal cells place fields

Previous patch-clamp recordings in PNs during spatial navigation identified key subthreshold features of activity within PFs, including increased theta power^40,43^ and an asymmetric ramp-like depolarization in subthreshold membrane potential (Vm)^12,40^. Notably, this Vm ramp asymmetry has been linked to an asymmetric plasticity kernel underlying behavioral time-scale synaptic plasticity (BTSP)^13^. Since little is known about the subthreshold features of spatial tuning in non-pyramidal cells, we sought to characterize these subthreshold signatures in SST place fields and compare them to our PN data.

Experimentally, both populations exhibited a large, slow depolarization within their PFs (Fig. 3b, Extended Data Fig. 3a), suggesting that, similar to PNs, SSTs’ place field formation might operate on a slow, seconds-long plasticity rule. However, only PNs showed the previously reported^40,43^ significant increase in theta amplitude within PFs (Fig. 3b,c; PNs: p<0.001, SSTs: p=0.6, paired t-test; Mean ± SEM = PNs, in-field: 0.l3 ± 0.006 spike heights, out-field: 0.l ± 0.004, SSTs, in-field: 0.16 ± 0.008, out-field: 0.l5 ± 0.008). Analysis of Vm ramp shapes revealed the expected asymmetric ramp in the PNs, which was driving a symmetric firing pattern inside the PF, similar to previous intracellular studies^13,40^ (Fig. 3d, Extended Data Fig. 3). Interestingly, the SST INs displayed a symmetric Vm ramp inside the place field (Fig. 3d,e), suggesting potential differences in the plasticity mechanism driving place field formation in SSTs compared with PNs. These differences may arise either from distinct post-synaptic mechanisms, or from differences in the structure of the inputs received by these populations. Specifically, CA1 PNs are assumed to receive relatively uniform spatial inputs from CA3^13,14^, whereas SSTs may integrate more spatially heterogeneous local inputs from CA1 PNs. Such differences may stem from subnetworks of PNs with overlapping or adjacent PFs converging onto the same SST+ interneuron, as proposed previously^44^.

**Fig. 3:**
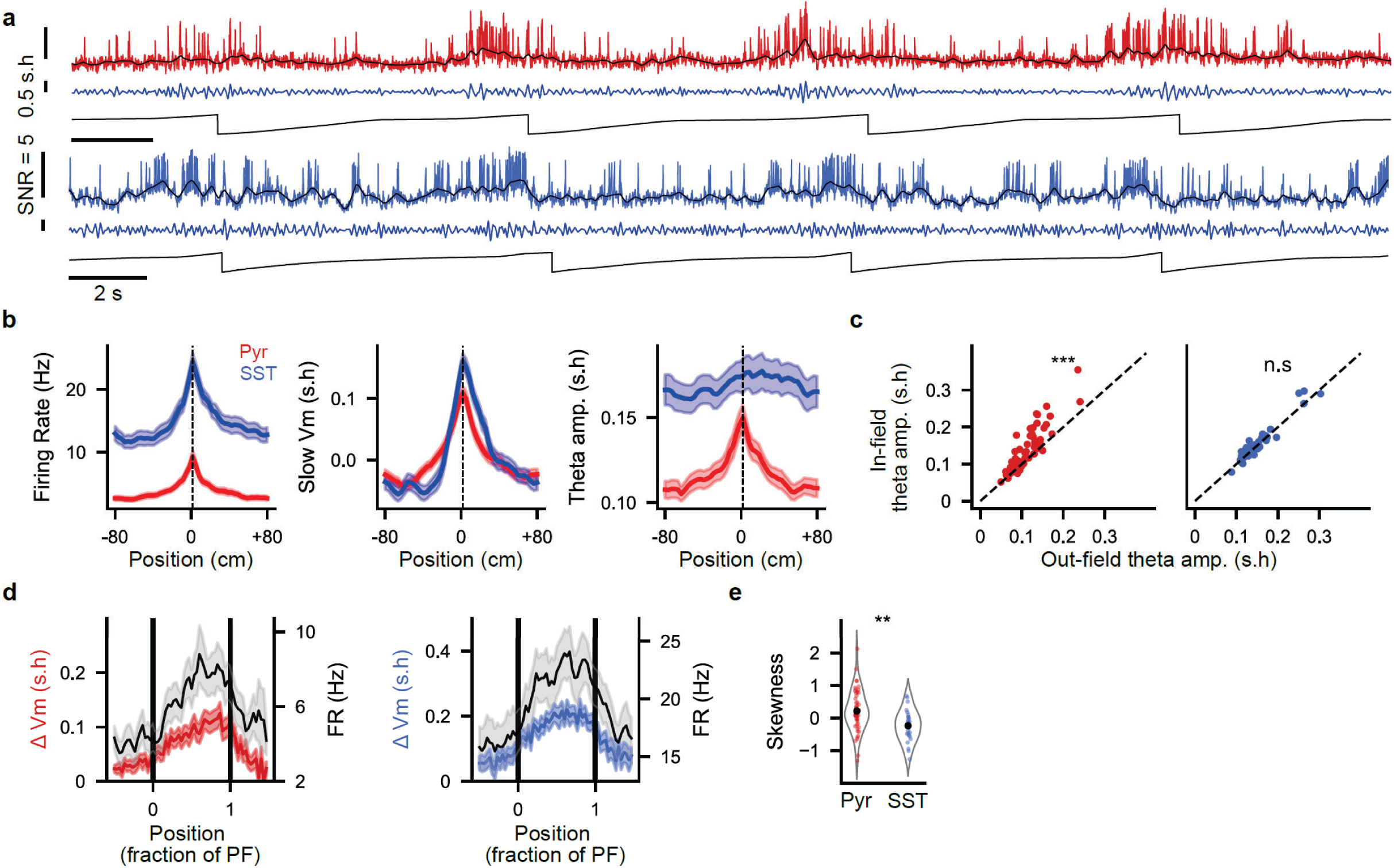
Subthreshold signatures of place field firing. **a**, Voltage traces a spatially tuned PN (top) and SST cell (bottom). Black: slow Vm signal (< 3 Hz), blue: theta-filtered signal (4-12 Hz). **b**, Population te (left), slow Vm (middle, s.h. - normalized to spike height), and theta amplitude (right) of all spatially tuned cells. For each cell, all re centered at the maximal value of the firing rate vector. **c**, In-field compared to out-field theta amplitude for all spatially tuned cells PNs, in-field: 0.13 ± 0.006, PNs, out-field: 0.1 ± 0.004, SSTs, in-field: 0.16 ± 0.008, SSTs, out-field: 0.15 ± 0.008. PNs: P = 1.6 X 10-6, paired t-test;). **d**, Population average shapes of slow Vm (colored) and firing rate (black) within the place field. Each place field y 50% to each side, and activity was re-binned into 60 spatial bins according to position within the place field. **e**, Skewness of the or all spatially tuned cells (P = 0.002, independent t-test).

### SST INs show non-spatial tuning properties

Previous calcium imaging studies demonstrated that several subtypes of interneurons, including SSTs, exhibit activity correlated with running speed^24–26^, consistent with their speed-modulated inputs from the medial septum^45^. However, since calcium imaging may not capture the temporal dynamics at high firing rates^19^ - such as those characteristic of SSTs (Fig. 1e,f) - we leveraged our access to their precise spiking activity and asked whether these cells display tuning beyond spatial selectivity. Agreeing with previous results, in addition to the relatively small subset of SSTs exhibiting spatial tuning (Fig. 2), many showed positive correlations with the animal’s running speed (Fig. 4c,d and Extended Data Fig. 4a,b). The firing rate-to-speed correlations varied on the single cell level (Extended Data Fig. 4b), and on average, speed-tuned SSTs increased their firing rate by ∼2.5 Hz per l0 cm/s (Fig. 4e). Notably, some cells were classified as both speed- and spatially tuned (Fig. 4f, 16 out of 83 speed-modulated interneurons). Overall, SSTs exhibited significantly stronger firing rate-speed correlations than PNs, with nearly 40% displaying significant speed tuning, compared to approximately 10% of PNs (Fig. 4g). Another large subset of SSTs fired uniformly across space, as determined using a spatial modulation test^24^ (Fig. 4b, 4g and Extended Data Fig. 4c). Uniform firing provides constant dendritic inhibition to PNs across the linear track, a pattern that may contribute to the enhancement of the spatial selectivity in PNs^46^.

**Fig. 4:**
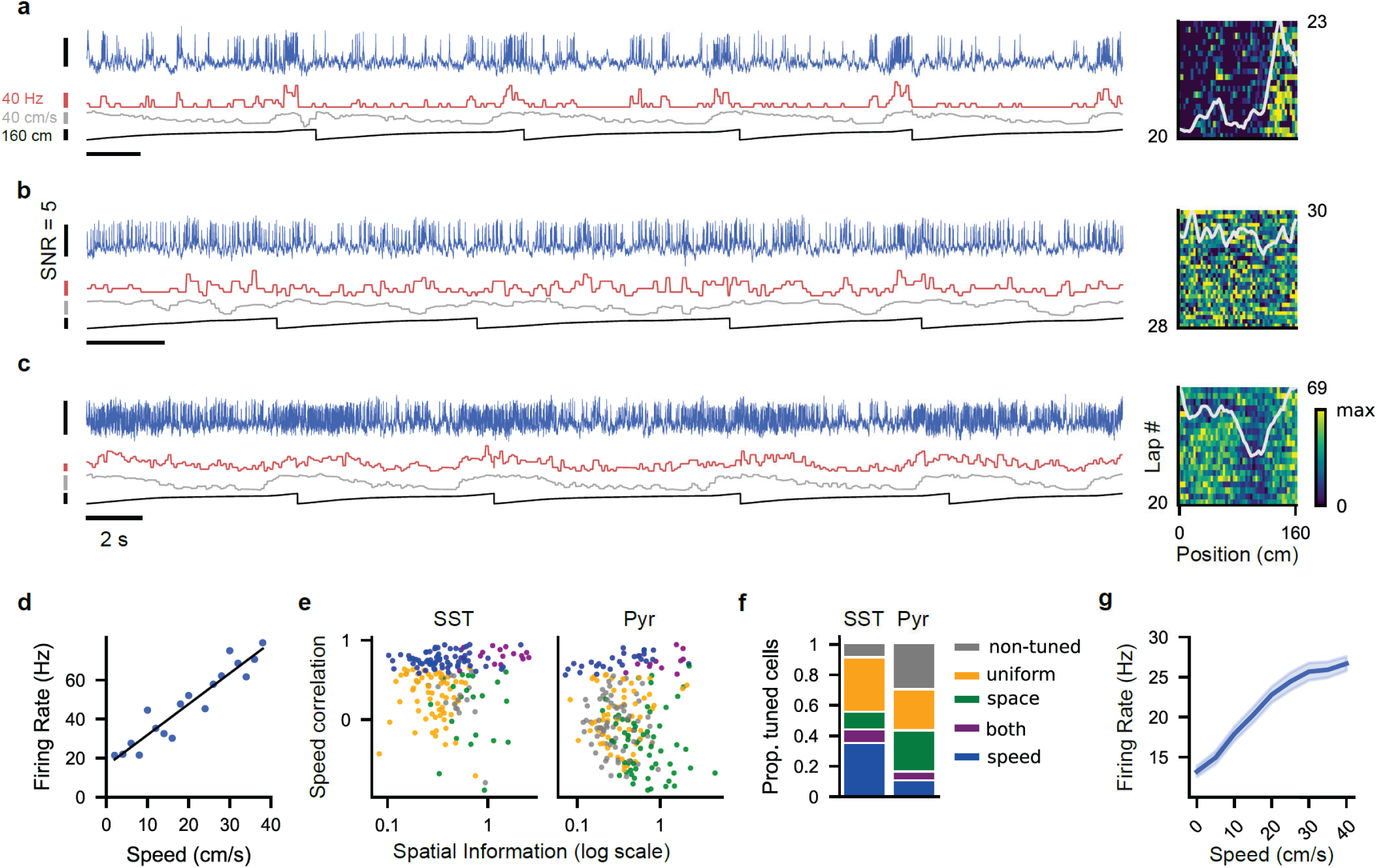
Non-spatial firing in SST+ interneurons. **a**, Left: Fluorescence trace (blue), firing rate (light red), running speed (gray), and position (black) for a spatially tuned SST+ interneuron across 5 laps. Right: Heatmap of firing rate by position for the same cell. Gray line indicates the average firing rate across all laps. Top right: maximal value of the average firing rate vector (Hz), bottom left: total number of laps in the trial. **b**, Same as **a** but for a uniformly firing SST+ cell. **c**, Same as **a** but for a speed-tuned SST+ cell. **d**, linear regression between firing rate and running speed for the speed-tuned cell shown in **c** (R = 0.92, P = 4.7 X 10-9). Speed was binned into 2 cm/s intervals. **e**, Scatter plot showing the Pearson R-value of the linear regression between firing rate and speed (y-axis) versus the spatial information index (x-axis, log scale) for all SSTs and PNs. Each point represents a cell, color-coded by its tuning type classification. **f**, Proportion of cells by tuning type within each neuronal population. **g**, Average firing rate by speed for all speed-tuned SST+ cells. Speed was binned into 5 cm/s intervals.

In summary, based on our access to their precise spiking dynamics, SST+ interneurons were classified into three tuning types: approximately 20% were spatially tuned, 40% were speed tuned, forming the largest subgroup, and 30% exhibited a spatially uniform firing pattern (Fig. 4g).

### Longitudinal recordings show unstable firing patterns in PNs and stable SST+ activity

Recent studies investigating the stability of spatial representations in CA1 over extended timescales have primarily focused on calcium imaging of PNs, showing varying levels of representational drifts^9–11,47,48^. However, calcium imaging lacks the temporal precision needed to resolve firing patterns, especially in interneurons, and cannot capture subthreshold dynamics. We thus sought to leverage the advantages of voltage imaging to longitudinally monitor both sub- and suprathreshold activity in PNs and SSTs across weeks, to study how the CA1 microcircuit balances stability and drift.

The CA1 pyramidal cell layer is extremely dense, and therefore, the reliable identification of the same PNs across weeks in longitudinal calcium imaging studies is challenging. Sparse expression of the voltage indicator was essential to ensure high SNR and allowed simple and reliable tracking of individual cells across sessions (Fig. 5a and Extended Data Fig. 5), despite limiting the experimental cell yield. We re-imaged the same FOVs ∼two weeks after the initial session and collected datasets of total 42 PNs and 44 SST+ cells, and quantified changes in their spiking and subthreshold properties.

**Fig. 5:**
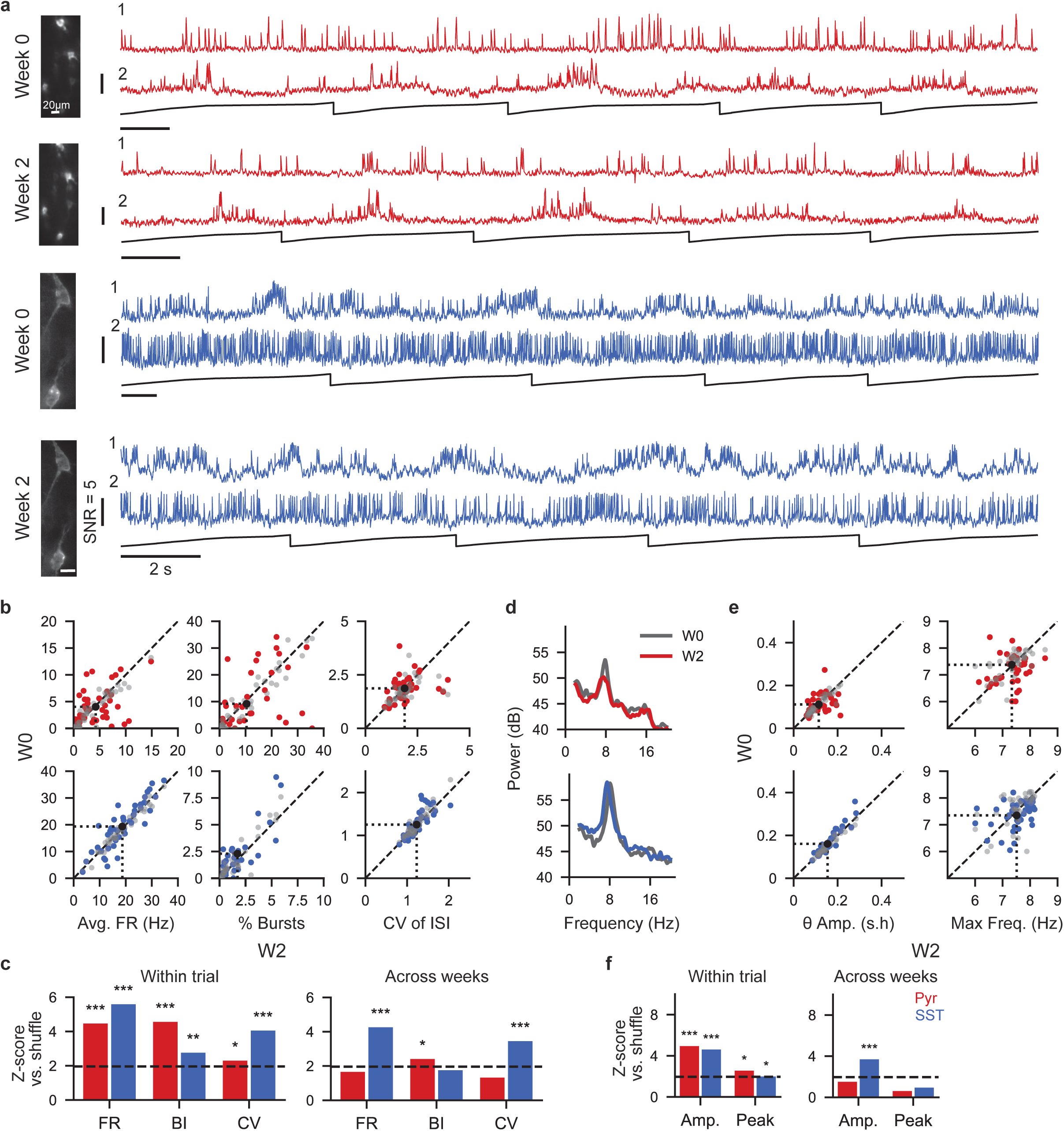
inter-population differences in stability of firing and subthreshold properties across weeks. **a**, Examples of the same PN (top) or SST (bottom) field of view recorded in two imaging sessions in a two-week interval. **b**, Average firing rate (left) percentage of spikes within bursts (middle) and inter-spike-interval (ISI) coefficient of variation (right) of all cells compared across two-weeks (13±2 days) (colored) or within two halves of the same trial as a control (gray). black dots and dashed lines denote mean population value for each week (N = 44 PNs and 42 SSTs). **c**, Left: Firing properties stability score within a trial in W0, first half compared to second half. Right: Same but for stability .across weeks (Z-scores of observed mean compared to shuffled means, * Z > 1.96, ** Z > 2.576, *** Z > 3.291, dashed line: Z = 1.96). **d** Example spectral densities of two cells recorded across weeks, one from each population, showing prominent theta peaks with slight differences in amplitudes and frequency components. **e**, Stability of subthreshold properties. Theta amplitude and peak frequency within the theta range were compared across weeks (colored) or within the same trial as a control (gray). **f**, Same as **c** but for subthreshold properties stability within a trial (left compared to across weeks (right).

For spiking metrics, we compared the average firing rate (FR), inter-spike-interval coefficient of variation (CV of ISI), and burstiness index (BI). At the population level, all metrics remained remarkably stable over time (Mean ± SEM = PNs; FR - W0: 4.23 ± 0.52, W2: 4.02 ± 0.52, BI - W0: 0.10 ± 0.02, W2: 0.09 ± 0.02, CV - W0: 1.88 ± 0.10, W2: 1.87 ± 0.09, SSTs; FR - W0: 18.68 ± 1.12, W2: 18.28 ± 1.12, BI-W0: 0.02 ± 0.002, W2: 0.02 ± 0.002, CV - W0: 1.23 ± 0.04, W2: 1.25 ± 0.05).

However, single-cell analysis revealed that PNs displayed substantial variability in FRs and CVs, and modest stability in their BIs (Fig. 5b, top and 5c). Notably, this stability was considerably lower than that observed within sessions (Fig. 5c, within session; FR: p<0.001, BI: p<0.001, CV: p=0.02, across weeks; FR: p=0.098, BI: p=0.014, CV: p=0.182, shuffled permutation test). In contrast, SST+ interneurons demonstrated remarkable stability in both FR and CVs of ISI, at a level comparable to within-session stability (Fig. 5b,c, within-session; FR: p<0.001, ISI CV: p<0.001, BI: p=0.007, across weeks; FR: p< 0.001, BI: p=0.092, CV: p<0.00l, shuffled permutation test).

Regarding subthreshold properties, SST+ interneurons maintained highly stable theta amplitudes across weeks, though their theta peak frequencies were less consistent (Fig. 5d-f, within session; theta amplitude: p<0.00l, peak frequency: p=0.053, across weeks; theta amplitude: p<0.001, peak frequency: p=0.346). In contrast, PNs showed reduced stability in both measures over time (within session; theta amplitude: p<0.00l, peak frequency: p=0.011, across weeks; theta amplitude: p=0.131, peak frequency: p=0.557).

These findings suggest that within the CA1 microcircuit, inhibitory activity patterns are more stable over time than excitatory ones. Specifically, the variability observed in PN firing across weeks is not shared with SSTs and therefore either originates from other local interneurons or excitatory inputs. The long-term stability exhibited by SSTs may result from integrating inputs across large populations of PNs or from additional stabilizing mechanisms - intrinsic or extrinsic – that balance their activity levels over time.

### Representational drifts in PNs and SSTs

A subset of the longitudinally recorded neurons from both populations was spatially tuned, and we next compared their tuning correlations across weeks to within-session controls (Fig. 6a). As apparent in the sorted heatmaps, PNs tuning was less reliable than SSTs when we compared two halves of the same imaging session (Fig. 6b), implying that PN representations are somewhat drifty even within the same imaging session, while SSTs are more consistent. Indeed, while the average within-session correlations of spatially tuned cells were similar between the cell types, the PNs’ correlations were significantly more variable (Fig. 6d, p=0.004, Levene test for equal variance, STDs=PNs: 0.33, SSTs: 0.l5), suggesting that PNs tuning is unstable even within the same imaging session, while SSTs are mostly stable. Of note, similar variability was observed when PNs were compared between odd and even sessions (Extended Data Fig. 6b), suggesting that this instability is not accumulated along the imaging session. Nevertheless, both populations showed larger center-of-mass distances across weeks, compared to within-session controls (Fig. 6c, PNs: p=0.01, SSTs: p =0.04, Wilcoxon signed-rank test. mean ± SEM = PNs: W0-W0: 6.99 ± 1.35, W0-W2: 12.41 ± 1.71, SSTs: W0-W0: 7.39 ± 1.50, W0-W2: 12.66 ± 2.69), indicative of a larger across-weeks representational drift. Similarly, both populations showed reduced correlations of their spatial tuning curves, though it was only significant for SSTs (Fig. 6d, PNs: p=0.13, SSTs: p = 6.6-3, paired t-test. mean ± SEM = PNs W0-W0: 0.43 ± 0.07, PNs W0-W2: 0.29 ± 0.07, SSTs W0-W0: 0.52 ± 0.04, SST control: 0.23 ± 0.07). However, for both cell types the population mean value was significantly larger than shuffled means (Fig. 6d, PNs; W0-W0: p<0.001, W0-W2: p=0.003, SSTs; W0-W0: p<0.001, W0-W2: p=0.004, Bootstrap permutation test), indicating that despite of apparent drift, significant spatial stability was maintained over the two-week interval.

**Fig. 6:**
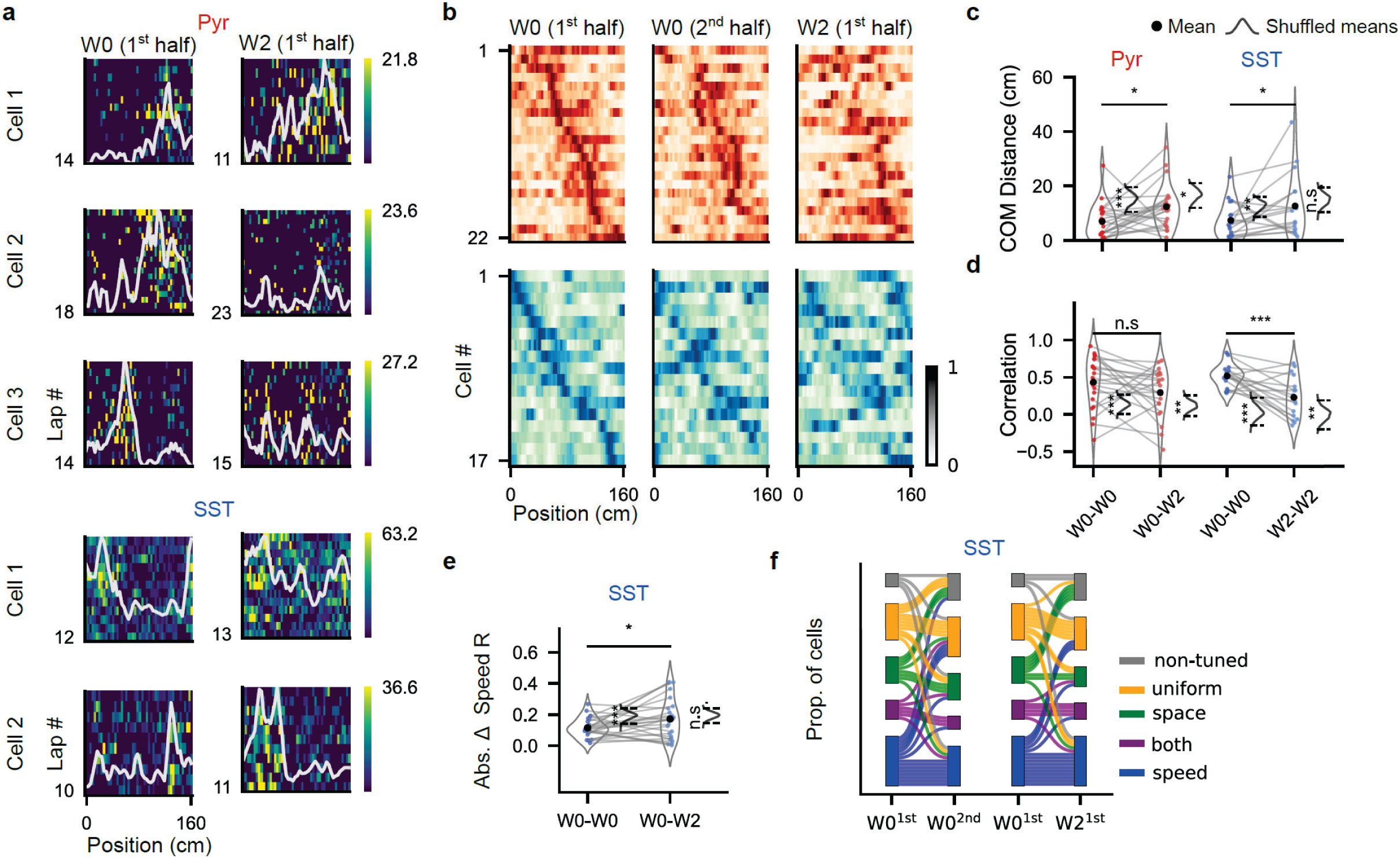
Drift in spatial and non-spatial tuning properties. **a**, Firing rate (FR) by space heatmaps from three PNs and two SSTs recorded across weeks. Average tuning curves (gray) are normalized by the maximal average FR value across both sessions. **b**, Average FR by position of all cells that were tuned in either W0 or W2, sorted by the position of their W0 place fields. W0 trials are split into two halves as a ‘W0-W0’ stability control. **c**, Center of mass (COM) distance compared within trial and across weeks (mean ± SEM = PNs: W0-W0: 6.99 ± 1.35, W0-W2: 12.41 ± 1.71, SSTs: W0-W0: 7.39 ± 1.50, W0-W2: 12.66 ± 2.69. PNs: P = 0.01, SSTs: P = 0.04, Wilcoxon signed-rank test). **d**, Tuning curve correlation between two halves of a trial in W0 compared to across weeks (mean ± SEM = PNs W0-W0: 0.43 ± 0.07, PNs W0-W2: 0.29 ± 0.07, SSTs W0-W0: 0.52 ± 0.04, SST control: 0.23 ± 0.07. PNs: P = 0.13, SSTs: P = 6.63 X 10-3, paired t-test). **e**, Absolute difference in FR-speed correlations compared within a trial and across weeks for all SST cells recorded in both weeks (mean ± SEM = W0: 0.115 ± 0.013, W2: 0.172 ± 0.026. p=0.03, paired t-test). **f**, Sankey diagrams for tuning type transitions across weeks.

As spatially tuned cells are a small subset of the SST population (Fig. 4g), we examined the stability of additional tuning types. Similar to spatial tuning, speed tuning was less stable across weeks compared to within-session controls, demonstrated by larger differences in FR-speed correlation values (Fig. 6e, p=0.03, paired t-test. mean ± SEM = W0: 0.115 ± 0.013, W2: 0.172 ± 0.026). Overall, for all three SST tuning classes, namely space, speed, and uniform, most cells retained their class identity while a fraction exhibited transition between the tuning types. Interestingly, the probability of transitioning between the tuning types was similar across weeks to within-session controls, indicating that beyond drift in their specific tuning properties, SST+ interneuron tuning types are at least partially maintained over time (Fig. 6f).

Taken together, the observed PN drift across weeks overall confirms previous calcium imaging studies^9–11,48^ at spiking resolution, but our data further reveal that PNs are noisy and unreliable both in their basic firing and subthreshold properties (Fig. 5) as well as in their spatial representations. Furthermore, our SST data suggest that drifts in inhibitory representations might be involved in the PN across-week drifts.

### Context switch induces Global remapping with opposite shifts of firing rates in both populations

Lastly, we examined the effects of context switching on both cell populations. To this end, we performed recording sessions that included abrupt transitions between environments, a manipulation shown to induce global remapping in CA1^27,49^. In each session, the animal performed the task in a familiar environment for 1.5 minutes before being teleported to a novel space with different visual cues but the same reward zone location and cue (Fig. 7a), where it continued the task for an additional 1.5 minutes. Mice exhibited premature licking and reduced speed in the novel environment, which gradually reverted with familiarity but persisted for over 5 exposures to this unfamiliar space (Fig. 7b and Extended Data Fig. 7a). Therefore, we analyzed only the first 5 remapping sessions of each animal.

**Fig. 7:**
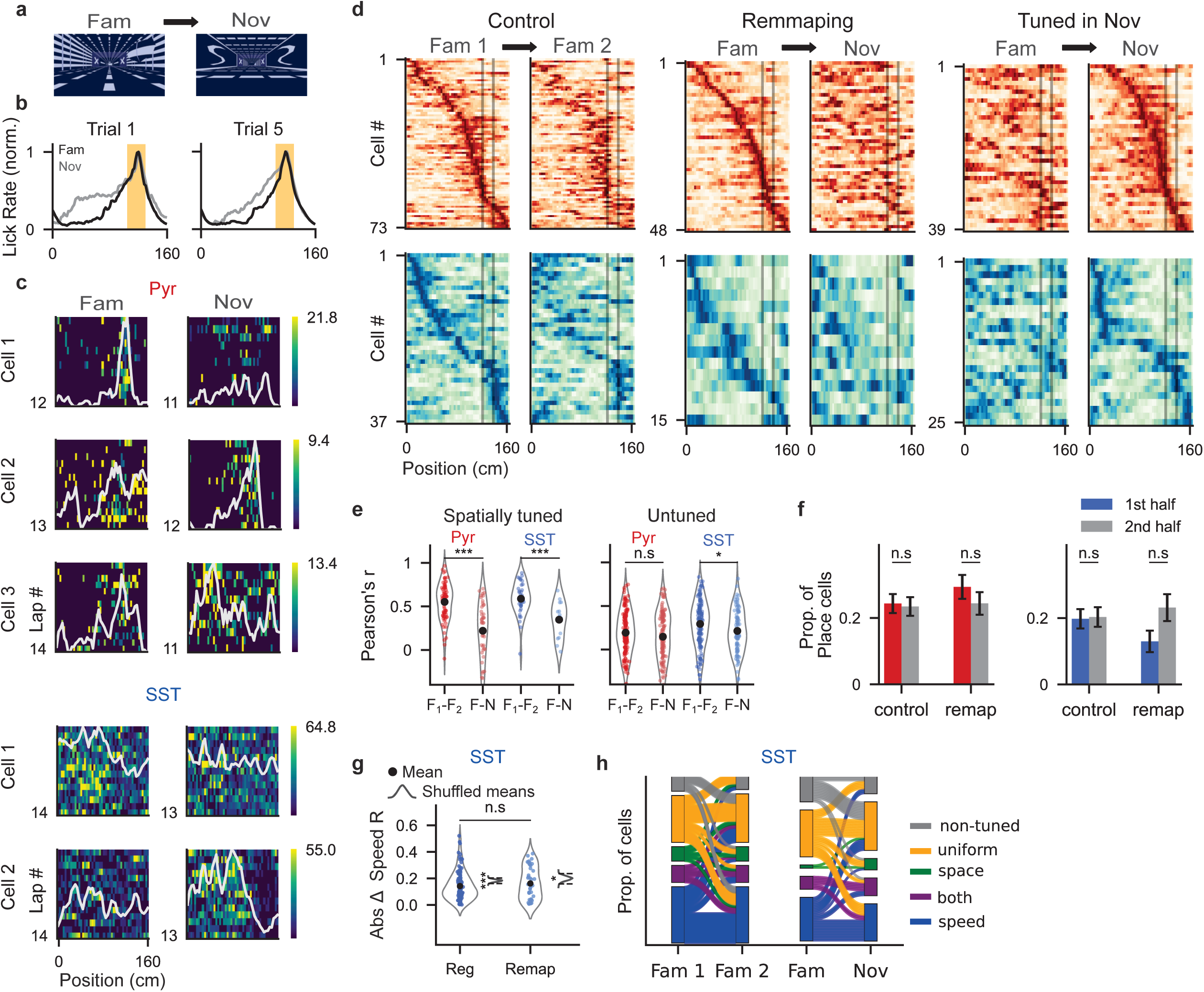
Global remapping in response to context switch. **a**, Top: familiar and novel virtual environments from the mouse’s point of view. **b**, Behavioral response to context switch, averaged across all mice. Mice show an increase in premature licking, a response that lasts for several exposures to the novel environment. **c**, Heatmaps of three PNs and two SSTs that alter their spatial activity following context switch from Fam to Nov context (gray line; average FR by position. **d**, Left: average FR by position of all spatially tuned cells in the familiar environment in week 0, sorted by max FR position in the first half. The first half of the trial is compared to the second half. Middle: same but for remapping trials in week 2, in which the second half of the trial is in the novel environment. Right: all cells that were spatially tuned in novel, by max FR position in novel. **e**, Left: mean correlation of firing rate vectors, either within (F1-F2) or across (F-N) contexts. Right: same for non-spatially tuned cells (Spatially tuned: PNs: P = 8.2 X 10^-10^, SSTs: P = 1.8 X 10^-3^, Untuned: PNs: P = 0.15, SSTs: P = 0.014, unpaired t-test). **f**, Probability of a cell being spatially tuned in each trial half (Control; PNs: P = 0.83, SSTs: P = 0.90, Remapping; PNs: P = 0.31, SSTs: P = 0.051, Two proportions Z-test). **g**, Absolute difference in FR-speed correlations across two halves of a trial in control and remapping experiments for all SST cells (mean ± SEM = Control: 0.142 ± 0.010, W2: 0.162 ± 0.018. P = 0.41, Wilcoxon rank-sum test.). **h**, Sankey diagrams for tuning type transitions across experiment halves in control and remapping experiments for all SST cells.

Exposure to a novel context induced remapping in spatially tuned cells of both populations, as indicated by a decreased correlation of spatial tuning curves compared to non-remapping controls (Fig. 7c-e). Even though the fraction of spatially tuned cells was not significantly different across environments, we observed an increase in the number of spatially tuned SST cells in the novel environment and a slight decrease in the tuned PNs (Fig. 7f, Control; PNs: p=0.83, SSTs: p=0.90, Remapping; PNs: p=0.3l, SSTs: p=0.05l, Two proportions Z-test), with more SST cells dedicated to the surroundings of the reward zone (Fig. 7d, right). Unlike the drift in speed-tuning that was observed across weeks (Fig. 6e), context switching had no effect on the consistency of FR-speed correlation values (Fig. 7g, p=0.41, Wilcoxon rank-sum test. mean ± SEM = Control: 0.142 ± 0.010, W2: 0.162 ± 0.018). Overall, similar to longitudinal experiments, SST tuning types showed similar transition probabilities as week 0 controls, displaying stability across environments (Fig. 7h)

In contrast with previous calcium imaging studies showing reduced SST calcium levels during context switch^24–28^, in our direct spiking recordings, most SST+ interneurons exhibited a global increase in firing rate following the transition, regardless of their tuning type. This increased firing activity gradually returned to baseline over about ∼7 laps. (Fig. 8a,b Extended Data Fig. 8a,b). In contrast, PNs showed a complementary decrease in firing rates (Fig. 8c,d and Extended Data Fig. 8d), which may be partially attributable to the increased SSTs activity. At the single-cell level, the proportion of SST+ interneurons with significantly increased firing (paired t-test, p < 0.05, comparing l0 laps before and after remapping) was tenfold higher compared to cells with significant decrease (22.5% vs. 2.5%, p < 0.001, Two proportions Z-test), an effect that was absent in controls (4.3% vs. 8.7%, p = 0.14). PNs showed the opposite trend, with a fourfold larger proportion of significantly decreasing cells (Remapping; 5% vs. 19%, p = 0.002, Control; 7% vs. 10%, p = 0.37). These shifts could not be explained by behavioral changes associated with novelty, such as reduced running speed, which would be expected to lower SSTs firing in speed-tuned cells (Fig. 8f). Instead, SST+ firing rates remained elevated across a range of running speeds in the novel environment (Fig. 8g).

**Fig. 8:**
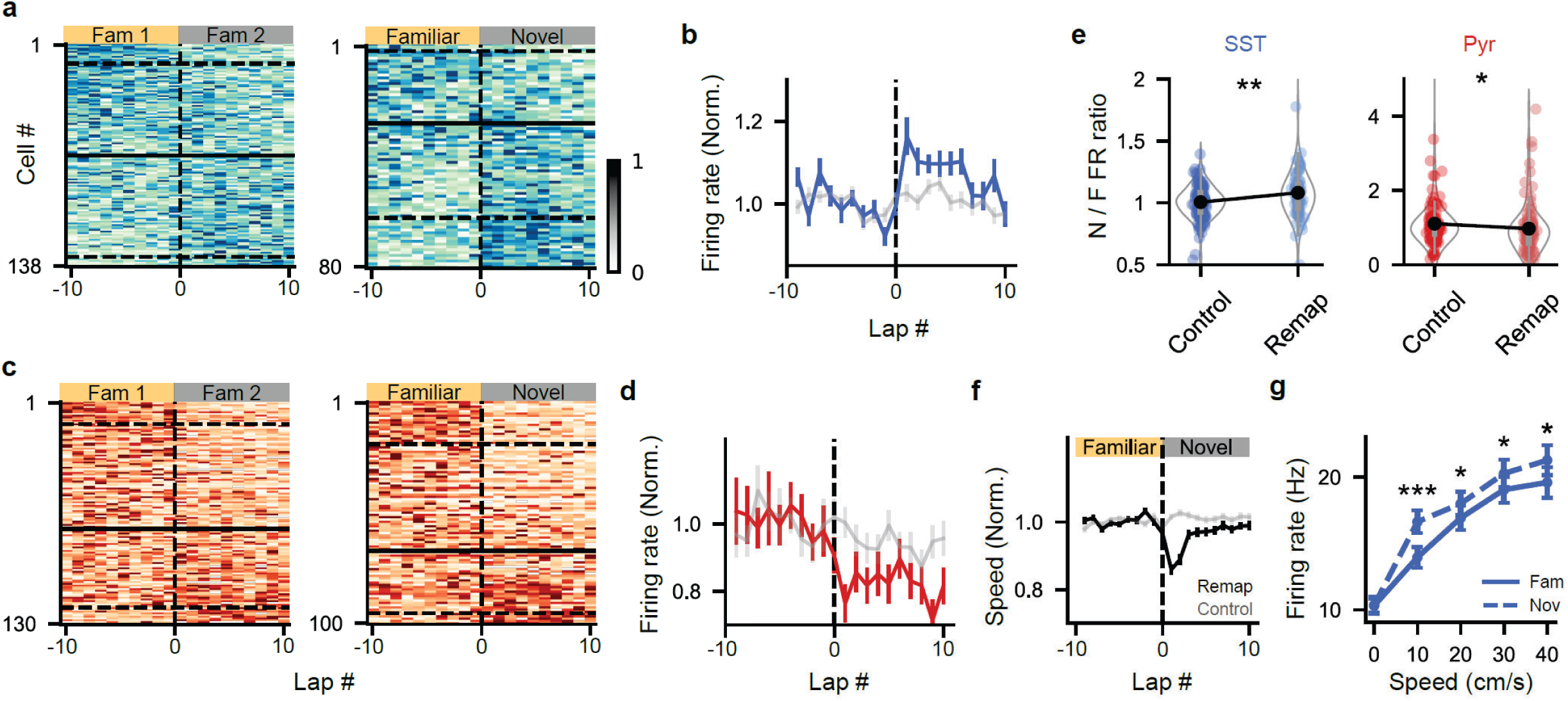
Opposite shifts of firing rates in SSTs and PNs during context switch. **a**, Firing rate (FR) by lap from ten laps before and after trial midpoint (dashed line) for all SST+ interneurons in control (Fam1-Fam2, left) and remapping (Fam-Nov, right) conditions. Each row was normalized from zero to one, and cells were sorted by their Nov/Fam FR ratio. Solid line marks the transition to Nov/Fam ratio >1, dashed lines mark cells with significant change in FR, either decrease (top line) or increase (bottom line). **b**, SST+ population average FR in remapping (blue) compared to control conditions (gray). For each condition, FR was normalized by the average FR across last ten laps before context switch (remapping) or trial midpoint (control). **c**, Same as **a** for all PNs. **d**, Same as **b** for all PNs. **e**, Violin plots of the FR ratio between novel and familiar environment for both populations (mean ± SEM = SST remapping: 1.08 ± 0.03, SST control: 1.00 ± 0.01, PN remapping: 0.97 ± 0.07, PN control: 1.11 ± 0.07. SSTs: P = 0.005, PNs: P = 0.009, Wilcoxon rank-sum test.). **f**, Average speed per lap across all trials from all mice, 10 laps before and after context switch (black) compared to control conditions (gray). For each trial, speed is normalized by the average speed in the familiar context. **g**, Firing rate in familiar (solid line) compared to novel (dashed line) across running speeds. Speed was binned into 10 cm/s bins (0-10 cm/s: P = 0.5,10-20: P = 2 X 10-5, 20-30: P = 0.01, 30-40: P = 0.02, 40-50: P = 0.033, Wilcoxon signed-rank test).

## Discussion

In this study, we used voltage imaging to obtain the first intracellular dataset from a large population of molecularly defined hippocampal neurons during both remapping and longitudinal experiments. By collecting data from SST+ cells and PNs under the same behavioral paradigm, we were able to directly compare their tuning properties, long-term stability, and responses to context switching. This approach recapitulated key features of PN place field activity previously identified through patch-clamp recordings, uncovered the distinct properties of SSTs dynamics and tuning, and confirmed previous calcium imaging studies showing representational drift of CA1 PNs. Our intracellular approach revealed that these drifts can be explained by the highly variable firing and subthreshold dynamics of PNs, and showed that while SST firing is much more reliable, their tuning properties do drift over a two-week period. Lastly, our data show that SST neurons increase their firing during context switch, suggesting that inhibitory control over PN dendrites might refine place cell formation and stabilization. These results provide mechanistic details into the stable and plastic properties of hippocampal circuit dynamics, and demonstrate that voltage imaging can extend previous findings to intracellular resolution, while scaling patch-clamp-like measurements to larger neuronal populations, complex behaviors, and additional cell types.

Many studies have compared spatial tuning between interneurons and PNs, yielding conflicting results regarding the differences in place field width and spatial information content^42,50,51^. In our dataset, we found no significant differences in either of the parameters. Notably, this is the first time SSTs’ spatial tuning has been assessed using intracellular recordings, measuring the subthreshold dynamics underlying their activity. Unlike PNs, which exhibited the expected asymmetric subthreshold ramps and increased intracellular theta amplitude around their place fields, SST+ cells displayed symmetrical ramps and no spatial modulation of theta amplitude.

Current models suggest that the asymmetric subthreshold ramp observed around PN place fields reflects an asymmetric plasticity kernel in time that potentiates spatially selective, uniformly distributed CA3 inputs. In contrast, SST+ interneuron place fields are unlikely to emerge through the same mechanism, given their reliance on both local PN-specific feedback and long-range excitatory inputs. Recently, there has been growing interest in understanding the plasticity mechanisms underlying interneuron spatial tuning and their implications on PN coding^52^. For example, Geiller et al. (2021) proposed that depression of excitatory-to-inhibitory (E–I) synapses following PN place field formation can induce inverse spatial tuning in their presynaptic interneurons. Whether similar mechanisms govern the emergence of positive interneuron place fields is still unclear. Nonetheless, our findings constrain the range of plausible plasticity mechanisms to those consistent with symmetrical subthreshold ramps and flat theta amplitude profiles. In addition, the absence of significant theta amplitude increases around interneuron place fields suggests that positive spatial tuning is either not directly inherited from PN place fields or that alternative theta sources dominate their subthreshold dynamics.

We found that ∼40% of SST+ interneurons were positively speed-tuned, in agreement with recent calcium imaging studies^24–26^ and extending their findings to intracellular resolution. Most of the remaining interneurons fired uniformly across space, suggesting that molecularly defined interneuron subtypes in CA1 can display heterogeneous functional roles. While spatially uniform inhibition has been proposed to enhance PN place fields by suppressing out-of-field excitation^46^, the functional role of speed-tuning in interneurons remains largely unknown. One possibility is that speed-modulated dendritic inhibition enables dynamic gating of entorhinal inputs based on locomotive state. Positive speed correlations have also been reported in PV+ interneurons^24,51^, which have been implicated in modulating phase precession in PNs^53,54^. Thus, speed signals within CA1 may influence both dendritic input selectivity and the fine temporal structure of spike timing within theta cycles, ultimately shaping memory encoding. However, direct evidence for these mechanisms, such as simultaneous recordings from place cells and their presynaptic speed-tuned interneurons, remains lacking.

Longitudinal intracellular studies using electrode-based techniques are essentially impossible, and therefore, precise spike-timing data were only feasible using extracellular techniques^34,55–57^. However, cell identity in these recordings is typically inferred from spike waveforms and activity measures, which have been shown to vary both across and within days, making long-term tracking highly error-prone^58^. Consequently, more recent studies extending over weeks and months have relied almost exclusively on calcium imaging. Yet calcium is a relatively slow proxy for neural activity, incapable of resolving individual spikes or reliably capturing the activity of high-firing neurons.

We demonstrate that voltage imaging enables repeated longitudinal recordings of both spiking and subthreshold dynamics across weeks, with reliable tracking of individual cells over time. Using this approach, we show that spatial representations of both PNs and SST+ cells exhibit drift over multi-week intervals. Our findings support the view that only a subset of PNs maintain place fields across weeks, but that sustained fields tend to be stable^9^. SST+ speed modulation was also less stable across weeks than within-session controls, but remained significantly correlated, extending prior findings of day-to-day speed tuning stability in VGAT-expressing interneurons^30^. While previous studies have reported similar proportions of speed-tuned cells across several interneuron subtypes^24^, it remains an open question whether their tuning stability over time is comparable to that of SSTs.

In terms of firing properties, we found that CA1 PNs exhibited substantially greater variability across weeks compared to SST+ interneurons. This finding aligns with prior work reporting greater drift in both spatial tuning and firing rates of CA1 versus CA3 PNs^33,41,59^, suggesting that CA1 is required for temporal coding^60^. Taken together, these observations suggest that the instability of CA1 PNs is not directly inherent from their CA3 or SST+ interneuron inputs, suggesting it is either resultant from changes in their synaptic weights or other potential sources of variability such as the entorhinal cortex.

An abrupt transition to a novel environment induced remapping of spatially tuned cells from both populations, consistent with recent calcium imaging reporting remapping across multiple interneuron subtypes, including SST+ cells^24^. Notably, those findings were obtained under different experimental conditions—a random foraging task on a conveyor belt with tactile cues—suggesting that remapping of spatially tuned interneurons is a general phenomenon that extends across sensory modalities and into goal-directed navigation in virtual reality environments. The observed decorrelation of spatial tuning curves was less pronounced in SST+ interneurons compared to PNs, consistent with previous findings showing spatial representation in CA1 interneurons rotates with visual cue rotation, but to a lesser extent than in PNs^30^. These results suggest that although interneuron spatial selectivity is context-dependent, it is more stable across contexts than that of PNs.

Remapping was accompanied by a transient increase in SST+ cells’ firing rates, regardless of their specific tuning profiles. This increase gradually returned to baseline over several laps and was mirrored by a decrease in PN firing rates. In contrast, several calcium imaging studies have reported reduced calcium signals in CA1 SST+ cells upon exposure to novel environments^24–28^. This discrepancy may stem from the variety of experimental setups, training paradigms or remapping protocols, all of which can influence animal behavior and responses to novelty. For example, studies examining PV+ interneuron responses to novel environments reported divergent results, including decreased^24,28^, increased^27^, or unchanged^25,48^ calcium activity. Importantly, to our knowledge, we are the first to report an increase in SST+ activity during remapping, despite experimental similarities to at least one prior study^27^. In that case, calcium signals were recorded from SST+ axons in stratum Oriens, likely originating from bistratified cells with somata in stratum Pyramidale. In contrast, we recorded SST+ somata in stratum Oriens, likely corresponding mostly to OLM cells^21^, which may account for the differing activity patterns observed.

As a potential circuit mechanism, OLM cells have been shown to receive direct excitatory input from cholinergic neurons in the medial septum^61^, which are active during attentive states and are thought to shift CA1 network dynamics towards memory encoding rather than consolidation^62,63^. Taken together, these findings suggest that in our experiments, cholinergic inputs from the medial septum may drive the firing of SST+ interneuron activity and suppress PN firing, reflecting a network state biased toward encoding in response to a novel context. This mechanism may support more complex interactions, as acetylcholine has also been shown to depolarize disinhibitory VIP+ basket cells^64^, pointing to a possible dual antagonistic influence on SST+ activity. Further studies using matched behavioral paradigms and complementary recording approaches are needed to resolve the discrepancies highlighted by our data.

Decreased dendritic inhibition is typically interpreted as facilitating dendritic plateau potentials and the formation of new place fields via BTSP. However, it is important to note that most CA1 PNs do not form new fields in each context. Thus, since increased SST+ cells activity was shown to reduce the size of the active PN ensemble^65^, we propose that dendrite-targeting interneurons, particularly OLM neurons targeting the PN apical tuft, rather than globally reducing their activity, may transiently increase it to promote the sparsening of PN population activity and enhance representational efficiency in novel environments.

Sheffield and Dombeck (20l9)^66^ proposed multiple dendritic mechanisms for place field formation, including both immediate and delayed onset fields, with or without engagement of BTSP. In our remapping paradigm, most place fields in the novel environment appeared on the first lap, and later emergence was typically not accompanied by prolonged depolarizations or other hallmarks of BTSP, although isolated examples were observed. Furthermore, we found no increase in the prevalence of complex spikes in novel environments, in agreement with previous work ^49^, suggesting that either burst firing did not play a major role in place field formation, or it did so in a small number of cells. Given that SST cell suppression has been associated with increased burst firing^53^, one might expect reduced bursting under the observed increase in SST activity. However, the reduction in PC firing rates in the novel environment was not specific to complex spikes (data not shown). This suggests that enhanced excitatory drive in the novel environment may be counterbalanced by increased dendritic inhibition, resulting in a net stabilization of burst firing relative to the familiar context.

## Methods

### Animals

Male and female mice aged 12–18 weeks, including 9 CKII-Cre (Jax #005359) and 5 SST-Cre (Jax #0l3044) mice, were used. All mice were heterozygous and maintained on C57BL/6 background. All animal experiments were conducted in accordance with guidelines approved by the Institutional Animal Care and Use Committee (IACUC) of the Hebrew University of Jerusalem.

### Surgery

Surgical procedures followed previously established protocols^31^. Mice were deeply anesthetized with 2% isoflurane, positioned in a stereotaxic frame, and maintained at ∼l.2% isoflurane throughout the procedure. Ophthalmic ointment was applied to prevent corneal drying, and body temperature was maintained at 37°C with a heating pad. The scalp was retracted, and the skull was cleaned and dried. A 3-mm craniotomy was performed over the left dorsal CA1 region (ML: 1.8 mm, AP: 2.3 mm) using a biopsy punch (Miltex). The dura was carefully removed with a 33G needle, and the cortex was aspirated under continuous saline irrigation to expose the external capsule. A central portion of the exposed axons was then gently removed to access the CA1 region.

For viral delivery, a microsyringe connected to a UMP3 microinjection pump (World Precision Instruments) was used to inject viral constructs at the center and four corners of the exposed CA1 surface. Each site was injected with 80 nl of virus at a rate of 2 nl/s at depth of 0.2 and 0.l mm (40 nl each). SST-Cre mice were injected with hSyn-DIO-somArchon1-P2A-somCheRiff-HA (final titer 9 × 10^12^ GC/ml). CKII-Cre mice were injected with a mixture of hSyn-fDio-somArchon-GFP (final titer 6 x l012 GC/ml) and EFla-DIO-FlpO (final titer l.72 x l010 GC/ml). All viral vectors were produced at the ELSC vector core facility (EVCF).

Following the injection, a stainless-steel cannula (2 mm length, 3 mm diameter; MicroGroup) was implanted and secured to the skull using 3M Vetbond tissue adhesive. Before implantation, the cannula was sealed with a 3 mm no. l round cover glass affixed with optical adhesive (NOA8l, Norland) and cured under UV light. A custom-made titanium head plate (eMachineShop) was then glued around the cannula, and any exposed skull was covered with dental cement (C&B Superbond, Sun Medical). The head plate included two threaded inserts to allow attachment of a plastic cap for protecting the glass from dirt. Animals were returned to their cages for recovery and received meloxicam (5 mg/kg) for two days.

### Behavioral Setup

Our custom virtual reality (VR) setup consisted of a 3D-printed wheel (l4 cm diameter) equipped with a rotary encoder (YUMO, 200 steps) mounted on the main axle and recorded rotations to control position in the VR environment in both forward and backward directions. The VR gain was adjusted so that the linear track spanned 160 cm while ensuring it did not correspond to a whole number of wheel rotations, preventing tactile or odor cues from providing task-relevant information.

The virtual environment was designed and rendered using ViRMEn (Virtual Reality MATLAB Engine)^67^ and projected onto a custom, compact 3D-printed screen. The projection was filtered with a 500 nm short-pass filter (Edmund Optics) to minimize spectral overlap with the near-infrared fluorescence of Archon. A lick port connected to a solenoid valve (Lee company LHD ported-style), was positioned in front of the animal, and licks were recorded using a capacitive touch sensor (SparkFun, AT42QTl0l0). For further information on the virtual reality and behavioral setup visit: https://github.com/yoavadamlab/VR-setup-for-NeuroImaging

### Optical setup

Imaging was performed on a Bergamo microscope equipped with a widefield epifluorescence module and a custom holography module (Thorlabs, described in^32^ In brief, Blue LED fluorescence was used to detect Archon-fused GFP tag for choosing FOVs. Archon imaging was done using a fiber-pigtailed 1 W 639 nm CW laser (CNI) patterned with a spatial light modulator (SLM, EXULUS-HD2, Thorlabs). The zero-order beam was blocked in an intermediate image plane using a glass slide with a deposited gold point.

Mice were imaged using a l6X 0.8 NA, 3-mm working distance objective (Nikon, Nl6XLWD-PF) and a custom 252 mm tube lens (effective magnification 20.lX). Laser intensity at the sample was 24 W/mm. Fluorescence was collected using a scientific CMOS camera (Hamamatsu, ORCA-Fusion) at 500 Hz.

### Behavioral Training and Longitudinal Imaging

Two weeks after surgery, following confirmation of viral expression and window clarity, mice underwent a 2–3-week behavioral training. Water was restricted to 1 mL per day, with free access to 2% citric acid water^68^ during weekends. Initially, mice were habituated to handling by experimenters, and when relatively calm, mice were placed on the wheel without virtual reality (VR) until they learned to rotate it efficiently (2–3 days, ∼20 minutes per day).

Training for the navigation task began with speed-dependent rewards, where mice received 4 μL of 10% sucrose water based on their running speed, with a progressively increasing speed threshold, with no VR. Mice were then trained in the familiar virtual environment for daily 40-minute sessions, receiving a reward upon entering the designated reward zone. After each lap, a 1-second black screen was presented before teleporting the mouse back to the starting position. Training initially followed an 8-minute “VR-on,” 2-minute “VR-off” protocol. Across sessions, the “VR-on” duration was gradually reduced to 4 minutes and then to 2 minutes, encouraging mice to accumulate rewards quickly once VR was active.

Once mice achieved a reward rate of >10 rewards per minute, imaging was conducted the following day. Imaging sessions lasted approximately 2 hours. Mice were first placed on the wheel without VR, and every time an imaging FOV was selected, a 3-minute VR session was initiated simultaneously with recording. The first recording session (Week 0) began by focusing on a distinct blood vessel junction, serving as the origin for this mouse’s “brain coordinates”. Coordinates were saved in ThorImage software (Thorlabs) together with the SLM masks of each recorded FOV. In the following sessions, we first re-centered on the same blood vessel junction, revisited the FOVs, and loaded the SLM masks to visually confirm cell positioning and identity across weeks. For remapping experiments, the context switch was initiated at the midpoint of each session. Once 1.5 minutes passed and a lap was completed, the environment switched to the novel one for the remainder of the session.

### Image segmentation and signal extraction

The signal extraction pipeline was adapted from the VolPy algorithm integrated into the CaImAn package (https://github.com/flatironinstitute/CaImAn)^69^. Briefly, movies were motion-corrected using the NoRMCorre algorithm^70^ with a kernel size of 3 for high-pass spatial filtering and rigid motion correction. Segmentation was performed based on the spatial light modulator (SLM) masks used for patterned illumination during recording.

A raw fluorescence trace was then obtained by averaging pixel intensities within the mask, and background noise was reduced using singular value decomposition (SVD) and ridge regression. The denoised trace was processed through an adaptive thresholding algorithm to identify the most prominent spikes. Extracted spike waveforms were averaged to generate a spike-shape template, which was then applied in a whitened-matched filter for re-detecting spikes. The resulting spike train was convolved with the template to generate a refined spiking trace.

To enhance signal localization, a weighted spatial footprint (SF) was estimated by ridge-regressing the spiking trace onto the high-pass filtered movie, assigning weights to pixels based on their contribution to the spiking signal. This SF was then applied to the segmented movie to extract a fluorescence trace with an improved signal-to-noise ratio (SF-trace). The spike detection and SF refinement process was repeated once more on the SF-trace, yielding an optimized fluorescence signal.

### Spike detection

For SST+ interneurons, spikes were extracted using the built-in spike detection algorithm in VolPy. Since PNs exhibit less uniform spike shapes due to bursting and complex spikes, the ridge regression by spike shapes did not perform as well. Therefore, we employed a simple thresholding approach for spike extraction - A high-pass filtered trace was obtained by subtracting a median filter (kernel size = 30 ms), and a spike detection threshold was defined as a specified number of standard deviations above the mean. To mitigate the effects of photobleaching, which reduces the signal-to-noise ratio (SNR) over time, we divided the fluorescence trace into two halves and performed spike detection separately for each.

To optimize this process, a custom graphical user interface (GUI) was developed to manually adjust the threshold and inspect the quality of detected spikes, ensuring accuracy in spike extraction.

### Subthreshold Vm Extraction and Normalization

To extract the subthreshold membrane potential (Vm), a median filter (kernel size = 30 ms) was applied to the SF-trace. Slow Vm components were obtained using a Butterworth low-pass filter (<3 Hz), while theta-range Vm was extracted with a band-pass Butterworth filter (4–12 Hz).

To compare subthreshold values and theta amplitudes across conditions, all subthreshold measures were normalized to the average spike amplitude within each 15-second segment of the fluorescence trace. Spike amplitude was measured locally as the difference between the peak fluorescence value and the lowest fluorescence value within the 6 ms preceding the peak. Mean theta amplitude was calculated as the average amplitude of the Hilbert-transformed theta-filtered trace.

### Spatial information, Spatial tuning, and place field detection

First, the linear track was divided into 50 spatial bins (∼3 cm each). The firing rate (FR) in each bin per lap was calculated as the number of spikes within that bin divided by the occupancy time. This generated an FR x space matrix, which was then averaged across all laps to obtain each cell’s spatial FR vector (spatial tuning curve).

Cells were classified as spatially tuned using a combined detection method^71^. First, a bootstrap test assessed the significance of each cell’s spatial information (SI) content, calculated in bits per second as detailed by Skaggs et al. (1992)^72^:

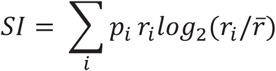

Where P*_i_* is the probability of the mouse occupying the *i^th^* spatial bin, r*_i_* is the firing rate within that bin, and 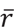 is the overall average firing rate.

To generate a shuffled SI distribution, we created l000 shuffled FR x space matrices by applying a random circular shift to each lap (row). A shuffled SI was calculated for each matrix, and a cell was considered spatially informative if its original SI exceeded 95% of shuffled values—this served as the first criterion.

Second, the cell’s in-field to out-field firing rate ratio had to exceed 2. Third, the cell was required to exhibit significant firing (Z>l.69 relative to all other bins in the FR matrix) in at least one bin within its place field (PF) for at least 30% of laps.

To define a cell’s PF, the spatial FR vector was smoothed using a three-point moving average. The PF was identified as all bins surrounding the peak of the spatial FR vector that exceeded a threshold of mean FR + 0.1(max FR – average FR). These bins determined the position and width of the PF.

### Analysis of Subthreshold Vm Ramps

For analyzing the shape for the slow Vm ramps accompanying PF firing, we followed previously described analysis^40^. Briefly, the Slow Vm trace and PF bins were computed as described above. To align all PFs regardless of their position and size, each PF was expanded by half of its original width in both directions. The expanded PF was then re-binned into 60 spatial bins. Then, the spatial slow Vm ramp was computed according to the new bins.

### Speed tuning

Speed-tuned cells were identified based on the correlation between firing rate and running speed. First, speed and firing rate were binned into l-second intervals. The time-binned speed vector was then further grouped into 20 speed bins, and the average firing rate within each bin was calculated. If a speed bin occupied less than 1% of frames it was removed from the analysis. A linear regression was performed between speed and firing rate, and cells were classified as speed-tuned if the regression yielded an r-value > 0.3 and a p-value < 0.01.

### Uniform firing

To classify a cell as spatially uniform, we applied a spatial modulation test for interneurons adapted from Geiller et al. (2020)^24^. First, l000 shuffled FR x space matrices were generated by randomly circularly shifting each row of the original FR x space matrix. Each shuffled matrix was then averaged across laps to generate a shuffled spatial FR vector.

For each spatial bin, the l0th and 90th percentile values were computed from the shuffled vectors and used as thresholds. A cell was classified as non-uniform if its original FR vector (smoothed with a window size of 3) exceeded either threshold for more than three consecutive spatial bins; otherwise, it was classified as uniform.

### Statistical tests

For each statistical test, we first assessed the normality of the data using the Shapiro-Wilk test. For paired samples, we applied a paired t-test if the differences were normally distributed, and a Wilcoxon signed-rank test otherwise. For non-paired comparisons, we used an independent t-test if the data in both groups were normally distributed, and a Wilcoxon rank-sum test otherwise. Proportions were compared using a two-proportions Z-test.

## Acknowledgments

We thank Noa Garty, Ron Buchnic, Mor Margolin, and Omer Cooper for their help with mouse training. We thank Maya Groysman and the ELSC vector core facility (EVCF) team for virus production, and Itamar Frachtenberg and the ELSC Fab Lab team for help with the design and fabrication of the VR behavioral setup. We thank Yaniv Ziv, Raunak Basu, and Yoram Burak for critically reading the manuscript. This work was supported by the European Research Council (ERC) starting grant #9487l6 and the Israel Science Foundation (ISF) grant #2940/24. YA is the Sachs Family Lecturer in Brain Sciences.

## Author contributions

RK performed surgeries, behavioral training, and *in vivo* imaging experiments. YM developed the VR setup and the signal extraction pipeline. RK analyzed all data with help from QY and YM. GS helped with mouse training and analyzed the behavioral data. SBS performed molecular biology and helped with surgeries and mouse training. RK and YA wrote the manuscript with input from all authors. YA provided funding and supervised all aspects of the project.

## Competing interests

The authors declare no competing interests.

---

**Extended Data Fig. 1:**
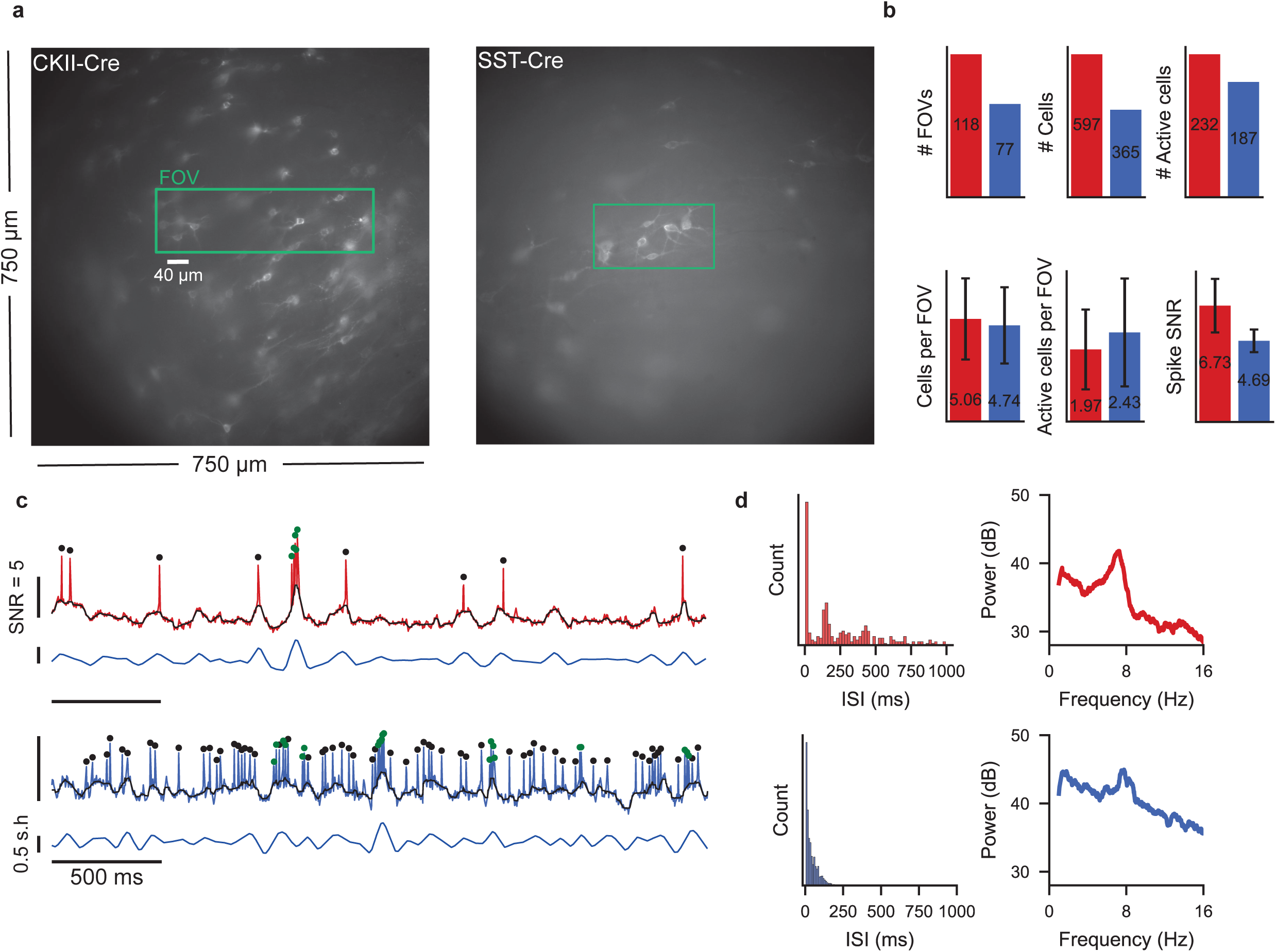
Recording statistics, spike detection and single cell examples. **a**, Example of a full frame image and a selected FOV for high-speed imaging in a CKII-Cre (left) and SST-Cre (right) mice. **b**, General recording statistics in week 0 recordings. Note that only active cells (cells with clear spikes and sufficient SNR - “Active cells”) were included in the analysis. **c**, Example fluorescence trace (colored), Median filtered trace (representing subthreshold Vm, black) and theta filtered signal (blue) for an example pyramidal neuron (top) and SST cell (bottom). simple spikes denoted in black, spikes within bursts in green. **d**, ISI distributions and power spectra for both cells in **c**.

**Extended Data Fig. 2:**
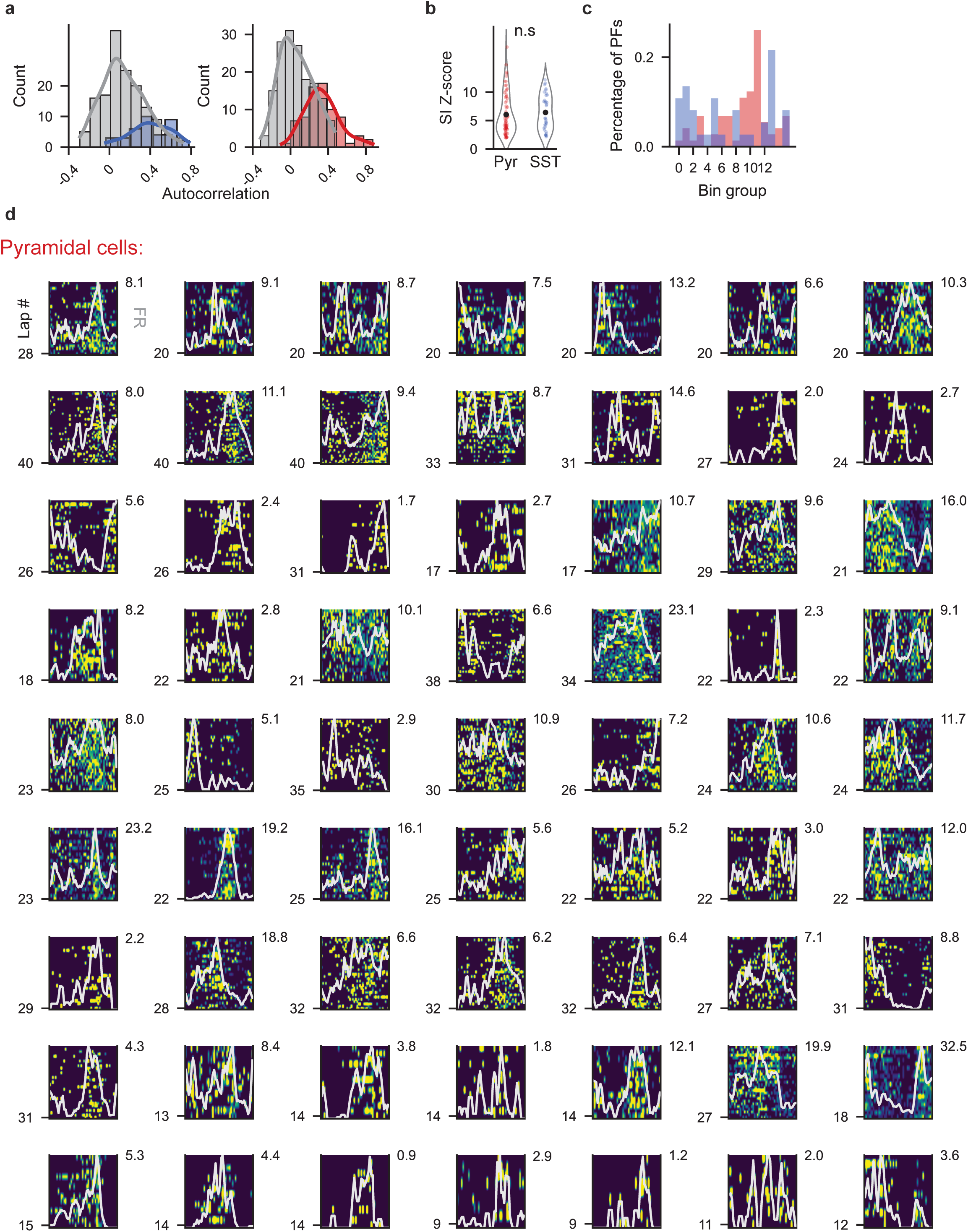
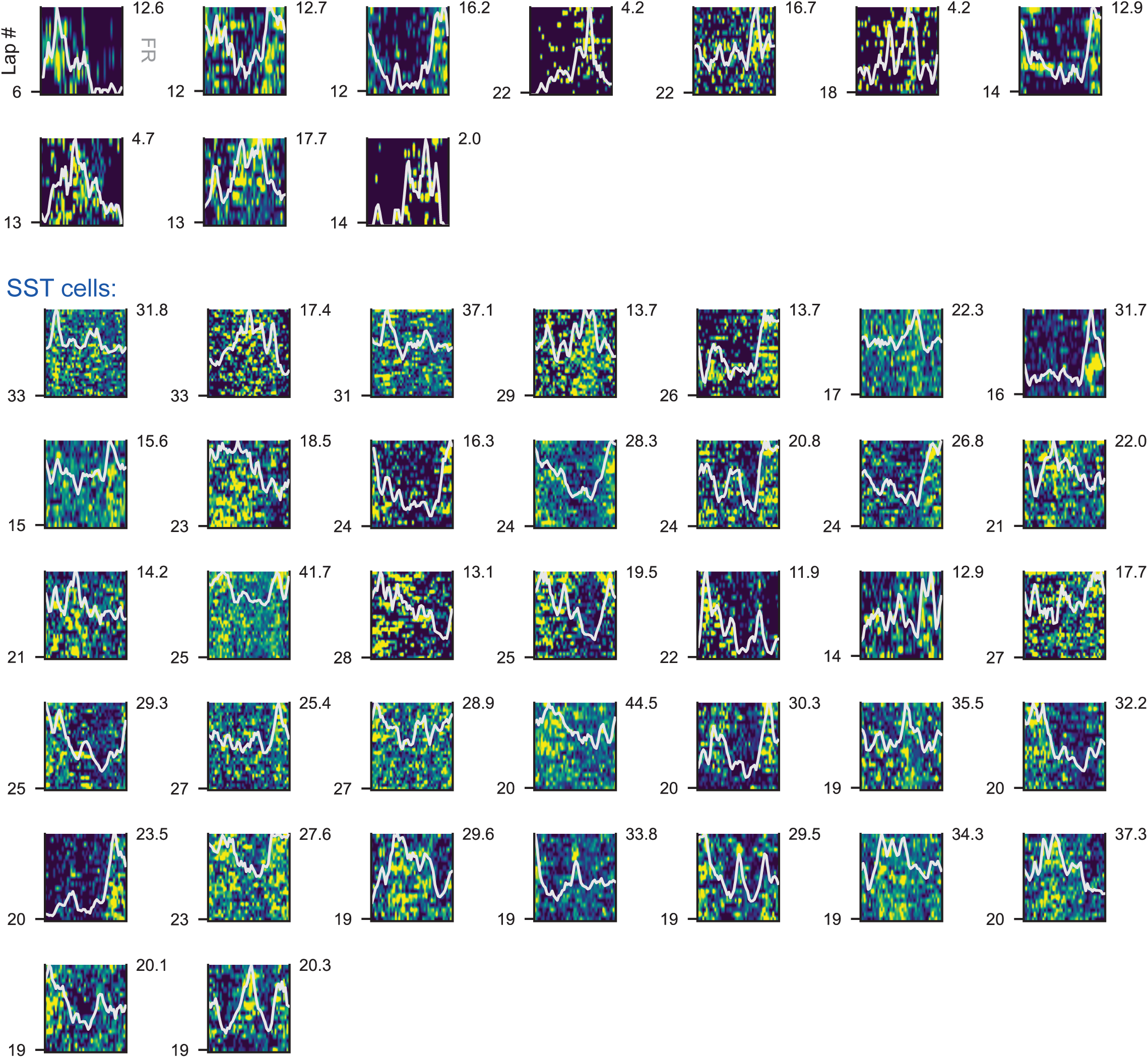
Additional spatial firing analysis and single cell firing rate X position heatmaps for all spatially tuned cells. **a**, Auto correlation of spatial firing rate (FR) vectors (1st half compared to 2nd half) of spatially tuned (colored) and non-tuned cells (gray) from each population. **b**, Z-scores of observed spatial information index compared to shuffled distribution for all spatially tuned cells. **c**, Fraction of place fields in each part of the track, showing PNs overrepresent the reward zone. Track position was binned into 16 10-cm bins. **d**, FR by position heatmaps of all spatially tuned cells. for each matrix, X axis is position, Y axis is number of laps (total is indicated at bottom left). Gray line denotes the spatial average FR vector (smoothed across 3 bins), its maximal value indicated at top right. Each color map limits are 0 to 1.5 times the maximal value.

**Extended Data Fig. 3:**
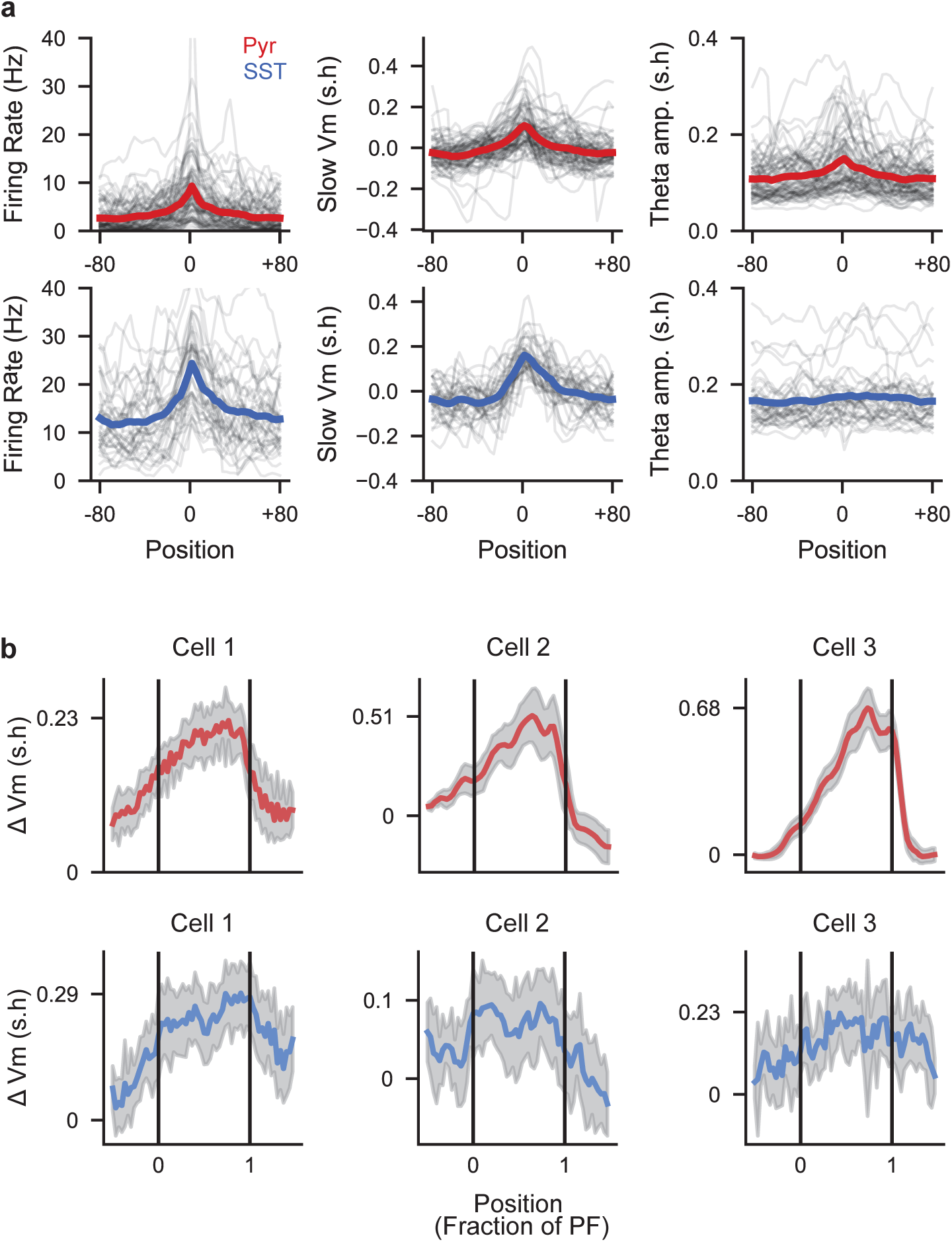
Spatial firing rate, slow Vm and theta amplitude. **a**, Firing rate (left), slow Vm (middle) and Theta amplitude (right) averaged across space for all spatially tuned cells, centered by the maximal value of the spatial firing rate vector. Individual cell average vectors are plotted in gray. **b**, Examples of slow Vm ramps within the place field of three spatially tuned cells from each population.

**Extended Data Fig. 4:**
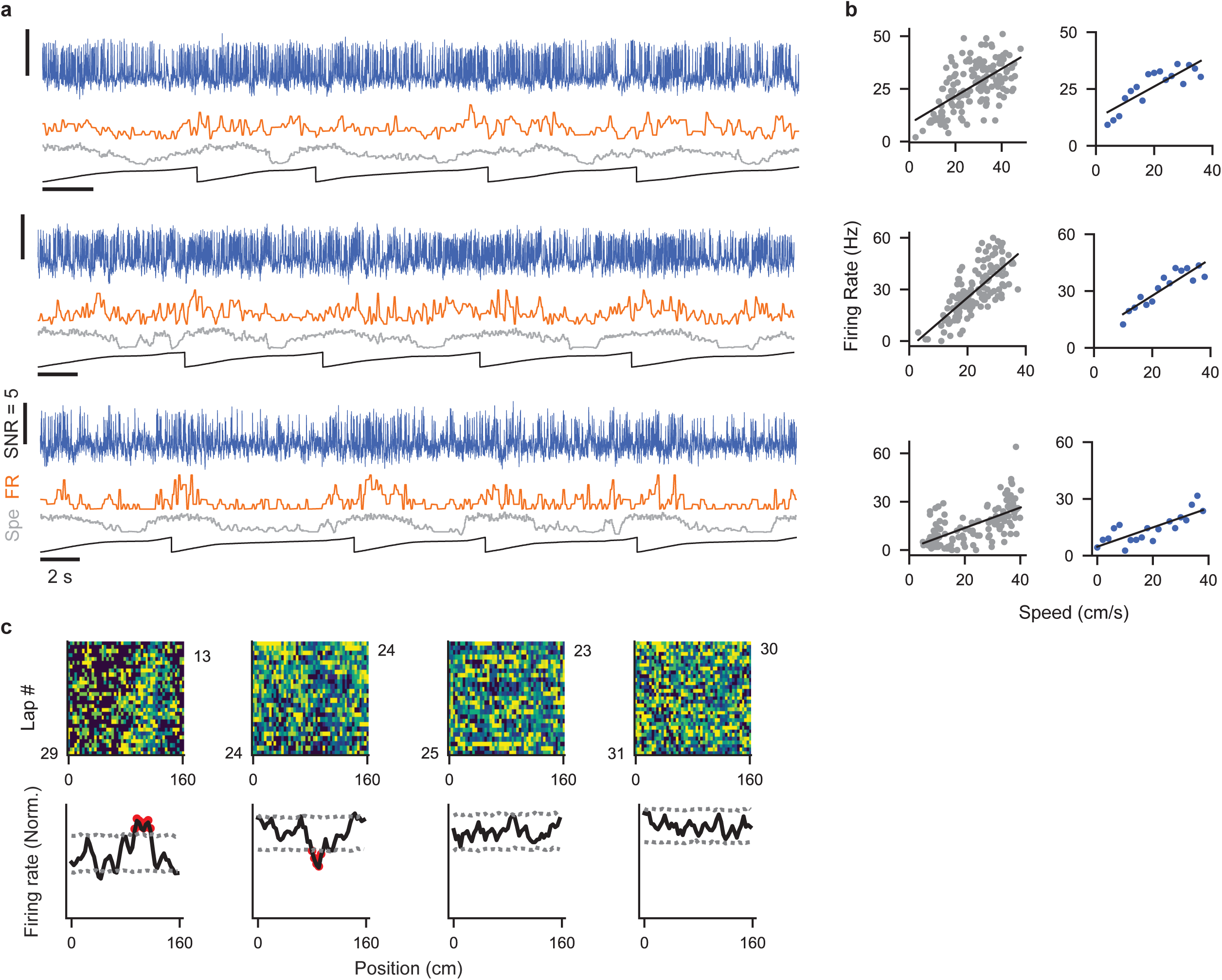
Speed-tuned SST cells and spatial uniformity test. **a**, Activity across 5 laps from three additional speed-tuned SSTs. **b**, Left - FR by velocity scatters for cells in **a**. each dot represents FR and average speed at each second of the trial. Right - Linear fit of FR averaged by speed bin. Speed is binned into 2 cm/s bins. **c**, Spatial uniformity test - to assess spatial uniformity of firing, each cell’s FR heatmap was shuffled 1000 times to calculate 5th & 95th percentiles of shuffled FRs in each spatial bin (gray dashed lines). If the FR exceeded these values for more than 5 consecutive bins, the cell was classified as spatially modulated. Uniform cells were defined as cells that were not spatially modulated or speed tuned.

**Extended Data Fig. 5:**
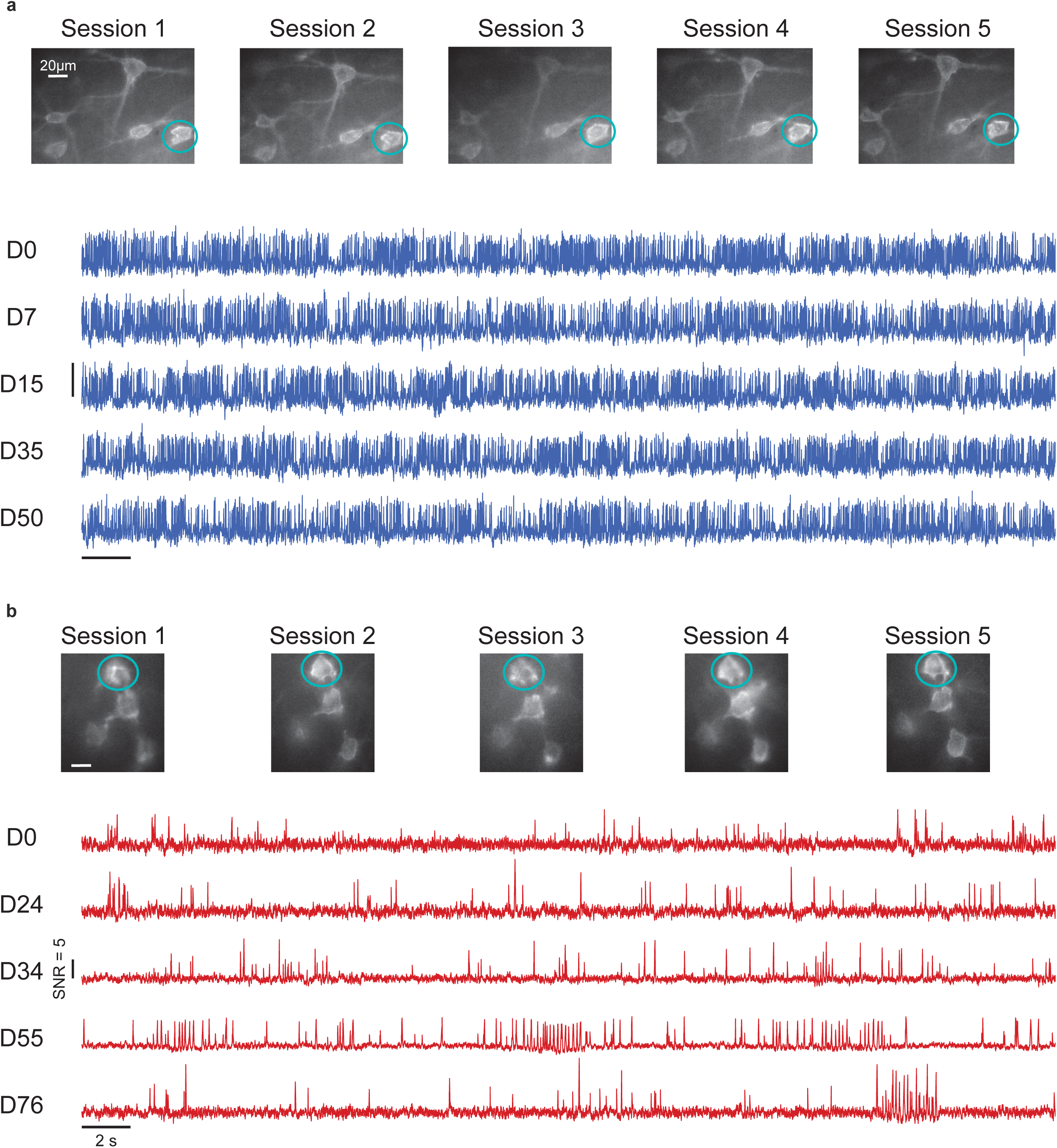
Longitudinal voltage imaging in multiple sessions over long time periods. **a**, Example of a field of view (FOV) of SSTs repeatedly imaged across five different sessions spanning more than 50 days and corresponding activity traces for the cell denoted by a circle. **b**, Same as **a** but for a FOV of pyramidal neurons.

**Extended Data Fig. 6:**
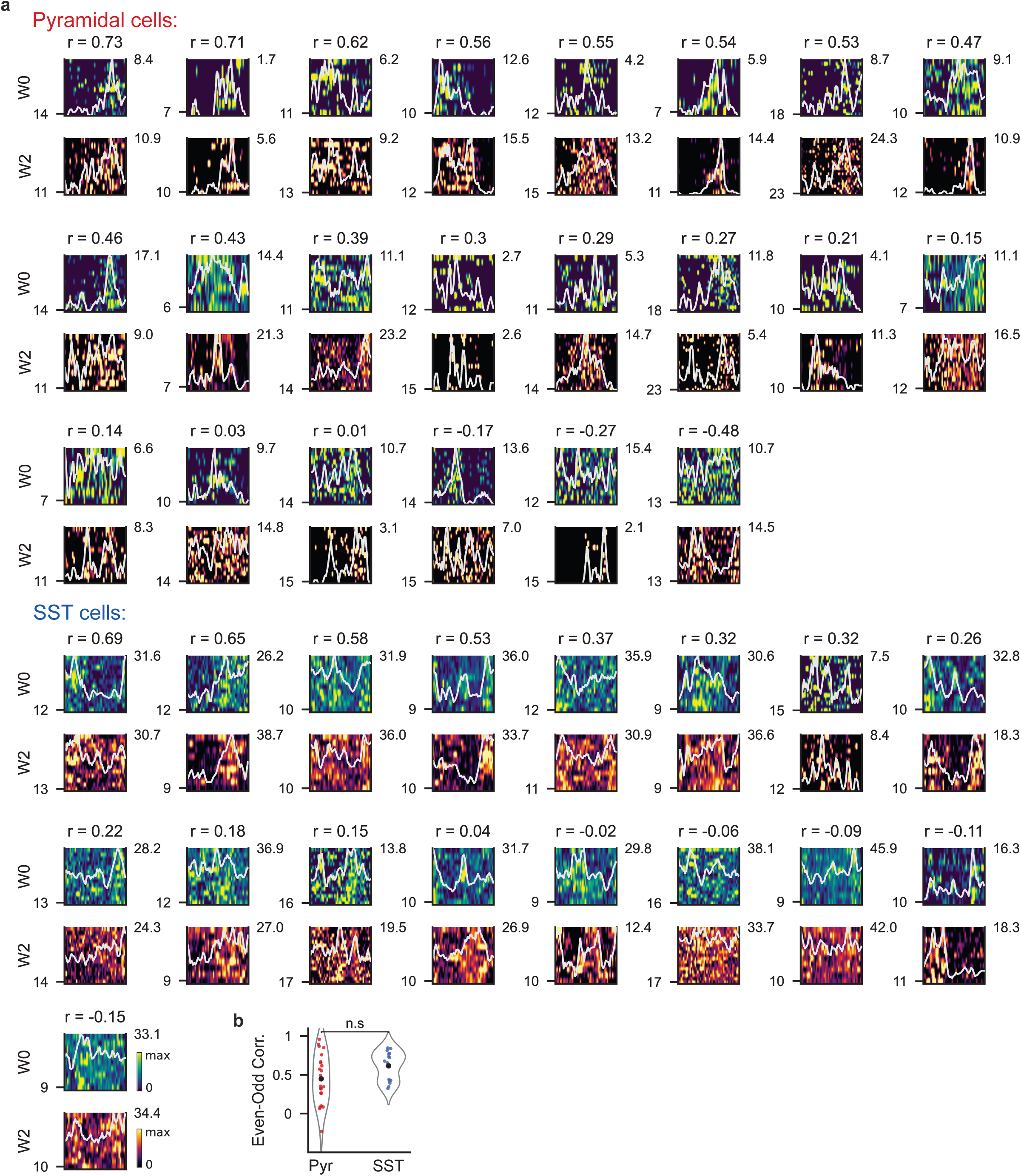
Firing rate X position heatmaps for longitudinally recorded spatially tuned cells. **a**, FR matrices of all cells that were spatially tuned in the first half of either W0 or W2 trials. W2 FR matrices are positioned below W0 matrices and plotted with a different firing rate colormap. Pairs were sorted by the cross-week correlation of their spatial tuning curves (gray lines). Correlation values are displayed above each pair. **b**, Related to Fig. 6D - correlation values between even and odd laps in control experiments show no-significant difference between populations but greater variability in PNs (mean ± STD = PNs: 0.45 ± 0.31, SSTs: 0.62 ± 0.18. P = 0.063, independent t-test. Leven’s test for equal variances: P = 0.015).

**Extended Data Fig. 7:**
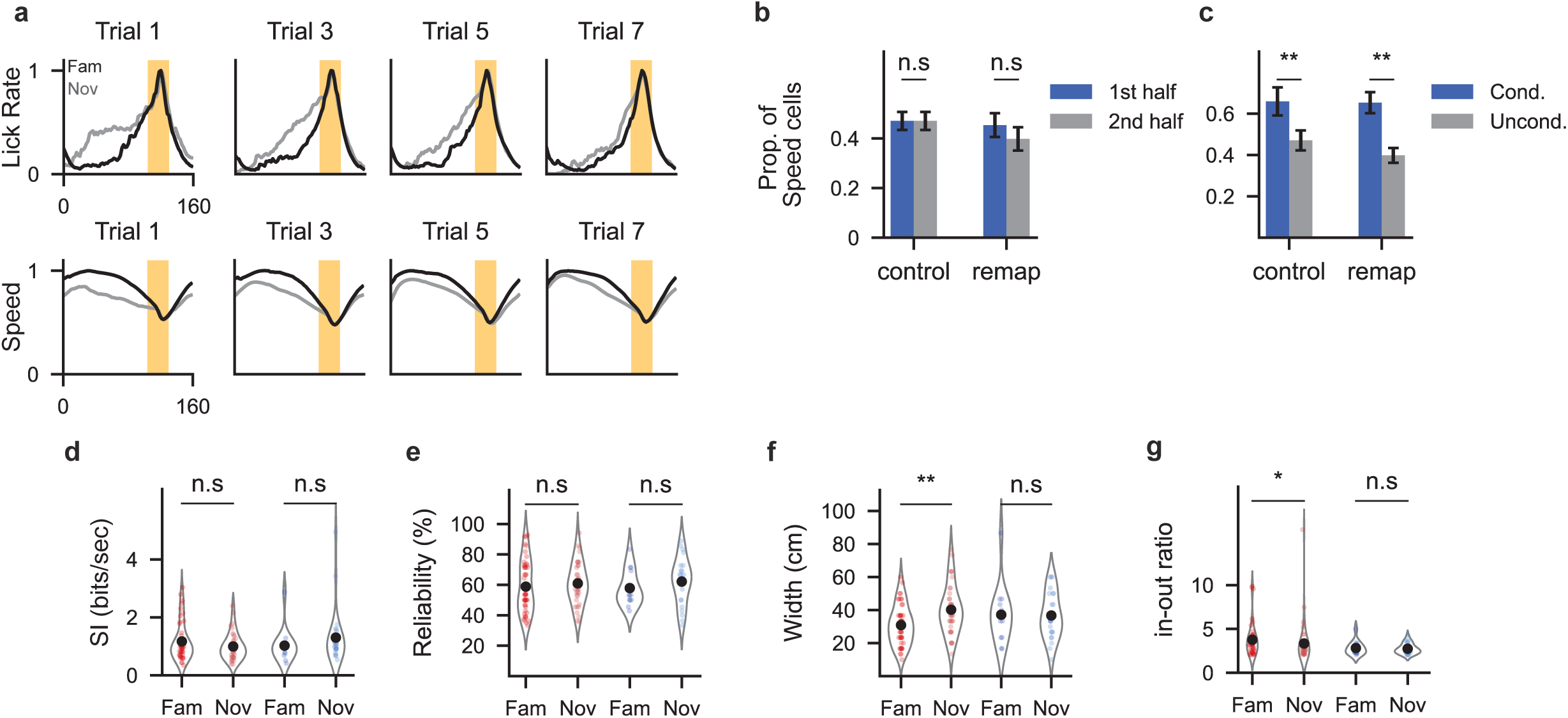
Effects of context switch on behavior, speed tuning and place field properties. **a**, Normalized lick rate (top) and speed (bottom) in familiar and novel environments in remapping trials 1,3,5 and 7, averaged across all mice. Upon exposure to novel, mice show premature licking and speed reduction, which gradually revert with familiarity. **b**, Proportion of speed-tuned SSTs in each half of a trial is unchanged in both remapping and control conditions (Control: P = 1, Remap: P = 0.41, Two proportions Z-test). **c**, The probability of an SST+ interneuron remaining speed tuned in the second half of a trial, given it was speed tuned in the first half (Cond) compared to unconditional probability (Uncond). Conditional probabilities were higher for both remapping and controls (Control: P = 0.003, Remap: P = 0.003, Two proportions Z-test). **d**, Comparison of Spatial information content across environments (PNs: P = 0.32, SSTs: P = 0.22, Wilcoxon rank-sum test). **e**, Percentage of laps with significant firing inside the place field (PF) (PNs: P = 0.36, SSTs: P = 0.32, Wilcoxon rank-sum test). **f**, PF widths (PNs: P = 0.003, SSTs: P = 0.89, Wilcoxon rank-sum test). g, In-field to out-field firing rate ratio (PNs: P = 0.02, SST: P = 1, Wilcoxon rank-sum test).

**Extended Data Fig. 8:**
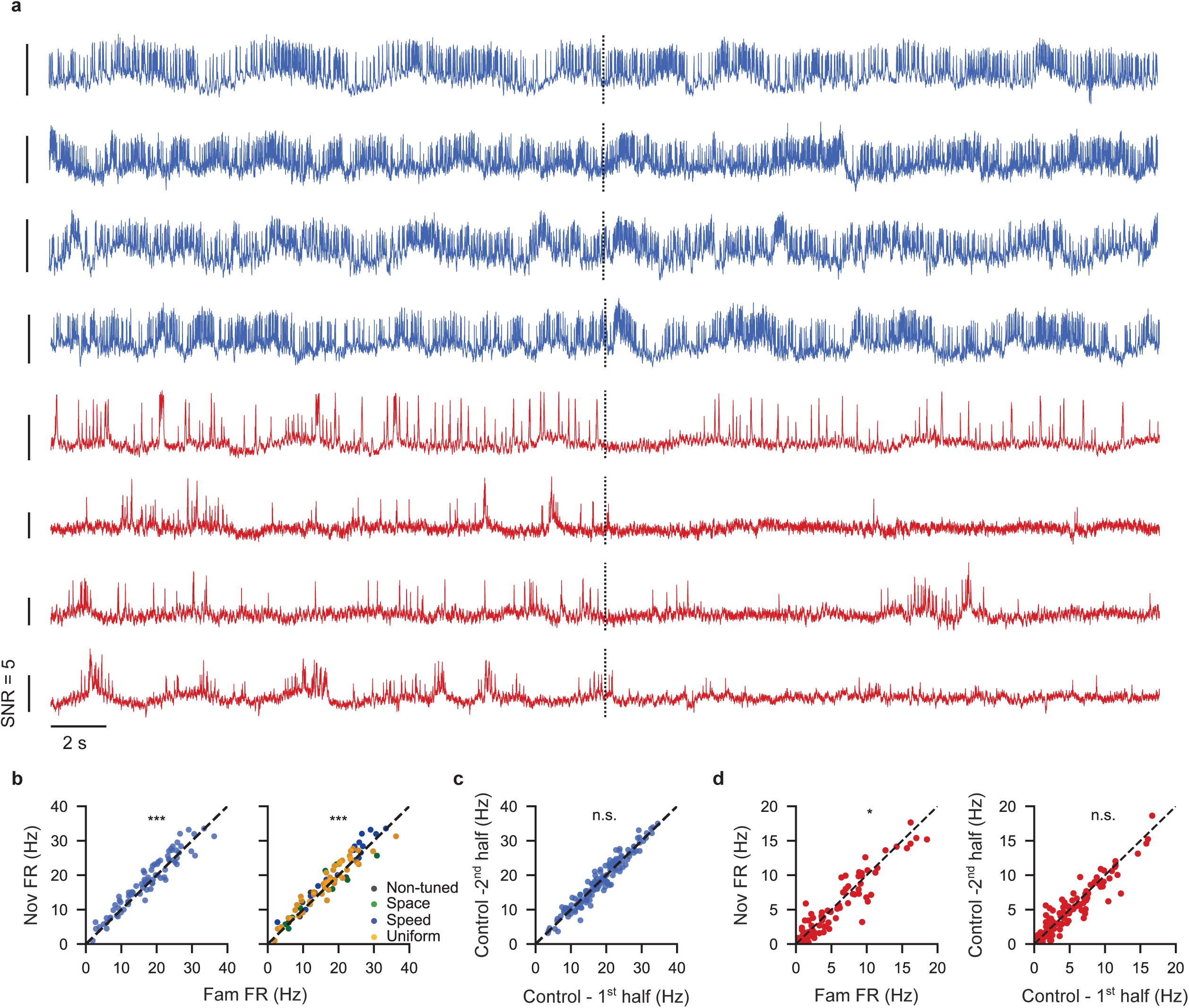
Cell-by-cell analysis of firing rate shifts during context switch. **a**, Example voltage traces of 10 seconds before and after context switch (dashed line) from 4 SST+ INs and 4 PNs demonstrating firing rate change following the switch. **b**, Average firing rate of the first 10 laps in the novel environment (Nov) compared to last 10 laps in the familiar environment (Fam). Right - same scatter where each dot is colored by the cells’ tuning type classification (mean ± SEM = Fam: 17.20 ± 0.87, Nov: 18.10 ± 0.87. P = 7.5 X 10^-3^, paired t-test). **c**, Same analysis but comparing two halves of an experiment with no remapping, to account for time passed (mean ± SEM = First half: 18.26 ± 0.62, Second half: 18.37 ± 0.64. P = 0.52, paired t-test.). **d**, Comparing average firing rates of PNs in remapping (left) and control (right) experiments (mean ± SEM = Fam: 4.79 ± 0.43, Nov: 4.47 ± 0.43, First half: 5.13 ± 0.41, Second half: 5.05 ± 0.41. Remap: P = 0.01, Control: P = 0.37, Wilcoxon signed-rank test).

## Notes

### Competing Interest Statement

The authors have declared no competing interest.

## References

1. O’Keefe, J., and Dostrovsky, J. (1971). The hippocampus as a spatial map: Preliminary evidence from unit activity in the freely-moving rat. Brain Research 34, 171–175. 10.1016/0006-8993(71)90358-1.

2. Burgess, N., Maguire, E.A., and O’Keefe, J. (2002). The Human Hippocampus and Spatial and Episodic Memory. Neuron 35, 625–641. 10.1016/S0896-6273(02)00830-9.

3. Moser, E.I., Kropff, E., and Moser, M.-B. (2008). Place Cells, Grid Cells, and the Brain’s Spatial Representation System. Annu. Rev. Neurosci. 31, 69–89. 10.1146/annurev.neuro.31.061307.090723.

4. Moser, M.-B., Rowland, D.C., and Moser, E.I. (2015). Place Cells, Grid Cells, and Memory. Cold Spring Harb Perspect Biol 7, a021808. 10.1101/cshperspect.a021808.

5. Ahmed, O.J., and Mehta, M.R. (2009). The Hippocampal Rate Code: Anatomy, Physiology and Theory. Trends Neurosci 32, 329–338. 10.1016/j.tins.2009.01.009.

6. Bostock, E., Muller, R.U., and Kubie, J.L. (1991). Experience-dependent modifications of hippocampal place cell firing. Hippocampus 1, 193–205. 10.1002/hipo.450010207.

7. Muller, R.U., and Kubie, J.L. (1987). The effects of changes in the environment on the spatial firing of hippocampal complex-spike cells. J. Neurosci. 7, 1951–1968. 10.1523/JNEUROSCI.07-07-01951.1987.

8. O’Keefe, J., and Conway, D.H. (1978). Hippocampal place units in the freely moving rat: Why they fire where they fire. Exp Brain Res 31, 573–590. 10.1007/BF00239813.

9. Ziv, Y., Burns, L.D., Cocker, E.D., Hamel, E.O., Ghosh, K.K., Kitch, L.J., Gamal, A.E., and Schnitzer, M.J. (2013). Long-term dynamics of CA1 hippocampal place codes. Nat Neurosci 16, 264–266. 10.1038/nn.3329.

10. Khatib, D., Ratzon, A., Sellevoll, M., Barak, O., Morris, G., and Derdikman, D. (2023). Active experience, not time, determines within-day representational drift in dorsal CA1. Neuron 111, 2348–2356.e4. 10.1016/j.neuron.2023.05.014.

11. Geva, N., Deitch, D., Rubin, A., and Ziv, Y. (2023). Time and experience differentially affect distinct aspects of hippocampal representational drift. Neuron 111, 2357–2366.e5. 10.1016/j.neuron.2023.05.005.

12. Bittner, K.C., Grienberger, C., Vaidya, S.P., Milstein, A.D., Macklin, J.J., Suh, J., Tonegawa, S., and Magee, J.C. (2015). Conjunctive input processing drives feature selectivity in hippocampal CA1 neurons. Nat Neurosci 18, 1133–1142. 10.1038/nn.4062.

13. Bittner, K.C., Milstein, A.D., Grienberger, C., Romani, S., and Magee, J.C. (2017). Behavioral time scale synaptic plasticity underlies CA1 place fields. Science 357, 1033–1036. 10.1126/science.aan3846.

14. Magee, J.C., and Grienberger, C. (2020). Synaptic Plasticity Forms and Functions. Annu Rev Neurosci 43, 95–117. 10.1146/annurev-neuro-090919-022842.

15. Fan, L.Z., Kim, D.K., Jennings, J.H., Tian, H., Wang, P.Y., Ramakrishnan, C., Randles, S., Sun, Y., Thadhani, E., Kim, Y.S., et al. (2023). All-optical physiology resolves a synaptic basis for behavioral timescale plasticity. Cell 186, 543–559.e19. 10.1016/j.cell.2022.12.035.

16. Buhl, E.H., Halasy, K., and Somogyi, P. (1994). Diverse sources of hippocampal unitary inhibitory postsynaptic potentials and the number of synaptic release sites. Nature 368, 823–828. 10.1038/368823a0.

17. Freund, T.F., and Buzsáki, G. (1996). Interneurons of the hippocampus. Hippocampus 6, 347–470. 10.1002/(SICI)1098-1063(1996)6:4<347::AID-HIPO1>3.0.CO;2-I.

18. Klausberger, T., and Somogyi, P. (2008). Neuronal Diversity and Temporal Dynamics: The Unity of Hippocampal Circuit Operations. Science 321, 53–57. 10.1126/science.1149381.

19. Klausberger, T., Magill, P.J., Márton, L.F., Roberts, J.D.B., Cobden, P.M., Buzsáki, G., and Somogyi, P. (2003). Brain-state- and cell-type-specific firing of hippocampal interneurons in vivo. Nature 421, 844–848. 10.1038/nature01374.

20. Lovett-Barron, M., Turi, G.F., Kaifosh, P., Lee, P.H., Bolze, F., Sun, X.-H., Nicoud, J.-F., Zemelman, B.V., Sternson, S.M., and Losonczy, A. (2012). Regulation of neuronal input transformations by tunable dendritic inhibition. Nat Neurosci 15, 423–430. 10.1038/nn.3024.

21. Müller, C., and Remy, S. (2014). Dendritic inhibition mediated by O-LM and bistratified interneurons in the hippocampus. Front. Synaptic Neurosci. 6. 10.3389/fnsyn.2014.00023.

22. Vaasjo, L.O., Kotermanski, S., Patel, T., Shi, H.J., Machold, R., and Chamberland, S. (2025). Dendritic inhibition terminates plateau potentials in CA1 pyramidal neurons. Preprint at bioRxiv, 10.1101/2025.06.05.657434 10.1101/2025.06.05.657434.

23. Klausberger, T., Márton, L.F., Baude, A., Roberts, J.D.B., Magill, P.J., and Somogyi, P. (2004). Spike timing of dendrite-targeting bistratified cells during hippocampal network oscillations in vivo. Nat Neurosci 7, 41–47. 10.1038/nn1159.

24. Geiller, T., Vancura, B., Terada, S., Troullinou, E., Chavlis, S., Tsagkatakis, G., Tsakalides, P., Ócsai, K., Poirazi, P., Rózsa, B.J., et al. (2020). Large-Scale 3D Two-Photon Imaging of Molecularly Identified CA1 Interneuron Dynamics in Behaving Mice. Neuron 108, 968–983.e9. 10.1016/j.neuron.2020.09.013.

25. Hainmueller, T., Cazala, A., Huang, L.-W., and Bartos, M. (2024). Subfield-specific interneuron circuits govern the hippocampal response to novelty in male mice. Nat Commun 15, 714. 10.1038/s41467-024-44882-3.

26. Udakis, M., Claydon, M., Zhu, H.W., and Mellor, J.R. (2024). Hippocampal OLM interneurons regulate CA1 place cell plasticity and remapping. Preprint, 10.1101/2024.11.11.622941 10.1101/2024.11.11.622941.

27. Sheffield, M.E.J., Adoff, M.D., and Dombeck, D.A. (2017). Increased Prevalence of Calcium Transients across the Dendritic Arbor during Place Field Formation. Neuron 96, 490–504.e5. 10.1016/j.neuron.2017.09.029.

28. Arriaga, M., and Han, E.B. (2019). Structured inhibitory activity dynamics in new virtual environments. eLife 8, e47611. 10.7554/eLife.47611.

29. Ali, F., and Kwan, A.C. (2019). Interpreting in vivo calcium signals from neuronal cell bodies, axons, and dendrites: a review. NPh 7, 011402. 10.1117/1.NPh.7.1.011402.

30. Schuette, P.J., Ikebara, J.M., Maesta-Pereira, S., Torossian, A., Sethi, E., Kihara, A.H., Kao, J.C., Reis, F.M.C.V., and Adhikari, A. (2022). GABAergic CA1 neurons are more stable following context changes than glutamatergic cells. Sci Rep 12, 10310. 10.1038/s41598-022-13799-6.

31. Adam, Y., Kim, J.J., Lou, S., Zhao, Y., Xie, M.E., Brinks, D., Wu, H., Mostajo-Radji, M.A., Kheifets, S., Parot, V., et al. (2019). Voltage imaging and optogenetics reveal behaviour-dependent changes in hippocampal dynamics. Nature 569, 413–417. 10.1038/s41586-019-1166-7.

32. Yang, Q., Baror-Sebban, S., Kipper, R., London, M., and Adam, Y. (2025). All-optical electrophysiology reveals behavior-dependent dynamics of excitation and inhibition in the hippocampus. Preprint at bioRxiv, 10.1101/2025.03.20.644347 10.1101/2025.03.20.644347.

33. Mankin, E.A., Sparks, F.T., Slayyeh, B., Sutherland, R.J., Leutgeb, S., and Leutgeb, J.K. (2012). Neuronal code for extended time in the hippocampus. Proc Natl Acad Sci U S A 109, 19462–19467. 10.1073/pnas.1214107109.

34. Muller, R.U., Kubie, J.L., and Ranck, J.B. (1987). Spatial firing patterns of hippocampal complex-spike cells in a fixed environment. J Neurosci 7, 1935–1950. 10.1523/JNEUROSCI.07-07-01935.1987.

35. Lever, C., Wills, T., Cacucci, F., Burgess, N., and O’Keefe, J. (2002). Long-term plasticity in hippocampal place-cell representation of environmental geometry. Nature 416, 90–94. 10.1038/416090a.

36. Thompson, L.T., and Best, P.J. (1990). Long-term stability of the place-field activity of single units recorded from the dorsal hippocampus of freely behaving rats. Brain Research 509, 299–308. 10.1016/0006-8993(90)90555-P.

37. Varga, C., Golshani, P., and Soltesz, I. (2012). Frequency-invariant temporal ordering of interneuronal discharges during hippocampal oscillations in awake mice. Proceedings of the National Academy of Sciences 109, E2726–E2734. 10.1073/pnas.1210929109.

38. Piatkevich, K.D., Bensussen, S., Tseng, H., Shroff, S.N., Lopez-Huerta, V.G., Park, D., Jung, E.E., Shemesh, O.A., Straub, C., Gritton, H.J., et al. (2019). Population imaging of neural activity in awake behaving mice. Nature 574, 413–417. 10.1038/s41586-019-1641-1.

39. Dombeck, D.A., Harvey, C.D., Tian, L., Looger, L.L., and Tank, D.W. (2010). Functional imaging of hippocampal place cells at cellular resolution during virtual navigation. Nat Neurosci 13, 1433–1440. 10.1038/nn.2648.

40. Harvey, C.D., Collman, F., Dombeck, D.A., and Tank, D.W. (2009). Intracellular dynamics of hippocampal place cells during virtual navigation. Nature 461, 941–946. 10.1038/nature08499.

41. Dong, C., Madar, A.D., and Sheffield, M.E.J. (2021). Distinct place cell dynamics in CA1 and CA3 encode experience in new environments. Nat Commun 12, 2977. 10.1038/s41467-021-23260-3.

42. Wilent, W.B., and Nitz, D.A. (2007). Discrete Place Fields of Hippocampal Formation Interneurons. Journal of Neurophysiology 97, 4152–4161. 10.1152/jn.01200.2006.

43. Epsztein, J., Brecht, M., and Lee, A.K. (2011). Intracellular Determinants of Hippocampal CA1 Place and Silent Cell Activity in a Novel Environment. Neuron 70, 109–120. 10.1016/j.neuron.2011.03.006.

44. Geiller, T., Sadeh, S., Rolotti, S.V., Blockus, H., Vancura, B., Negrean, A., Murray, A.J., Rózsa, B., Polleux, F., Clopath, C., et al. (2022). Local circuit amplification of spatial selectivity in the hippocampus. Nature 601, 105–109. 10.1038/s41586-021-04169-9.

45. Fuhrmann, F., Justus, D., Sosulina, L., Kaneko, H., Beutel, T., Friedrichs, D., Schoch, S., Schwarz, M.K., Fuhrmann, M., and Remy, S. (2015). Locomotion, Theta Oscillations, and the Speed-Correlated Firing of Hippocampal Neurons Are Controlled by a Medial Septal Glutamatergic Circuit. Neuron 86, 1253–1264. 10.1016/j.neuron.2015.05.001.

46. Grienberger, C., Milstein, A.D., Bittner, K.C., Romani, S., and Magee, J.C. (2017). Inhibitory suppression of heterogeneously tuned excitation enhances spatial coding in CA1 place cells. Nat Neurosci 20, 417–426. 10.1038/nn.4486.

47. Clopath, C., Bonhoeffer, T., Hubener, M., and Rose, T. (2017). Variance and invariance of neuronal long-term representations. Philosophical Transactions of the Royal Society B: Biological Sciences 372, 20160161. 10.1098/rstb.2016.0161.

48. Hainmueller, T., and Bartos, M. (2018). Parallel emergence of stable and dynamic memory engrams in the hippocampus. Nature 558, 292–296. 10.1038/s41586-018-0191-2.

49. Cohen, J.D., Bolstad, M., and Lee, A.K. (2017). Experience-dependent shaping of hippocampal CA1 intracellular activity in novel and familiar environments. eLife 6, e23040. 10.7554/eLife.23040.

50. Ego-Stengel, V., and Wilson, M.A. (2007). Spatial selectivity and theta phase precession in CA1 interneurons. Hippocampus 17, 161–174. 10.1002/hipo.20253.

51. Arriaga, M., and Han, E.B. (2017). Dedicated Hippocampal Inhibitory Networks for Locomotion and Immobility. J. Neurosci. 37, 9222–9238. 10.1523/JNEUROSCI.1076-17.2017.

52. Jeong, N., and Singer, A.C. (2022). Learning from inhibition: Functional roles of hippocampal CA1 inhibition in spatial learning and memory. Current Opinion in Neurobiology 76, 102604. 10.1016/j.conb.2022.102604.

53. Royer, S., Zemelman, B.V., Losonczy, A., Kim, J., Chance, F., Magee, J.C., and Buzsáki, G. (2012). Control of timing, rate and bursts of hippocampal place cells by dendritic and somatic inhibition. Nat Neurosci 15, 769–775. 10.1038/nn.3077.

54. Chadwick, A., van Rossum, M., and Nolan, M. (2015). Modelling phase precession in the hippocampus. BMC Neurosci 16, O18. 10.1186/1471-2202-16-S1-O18.

55. Kentros, C.G., Agnihotri, N.T., Streater, S., Hawkins, R.D., and Kandel, E.R. (2004). Increased Attention to Spatial Context Increases Both Place Field Stability and Spatial Memory. Neuron 42, 283–295. 10.1016/S0896-6273(04)00192-8.

56. Kentros, C., Hargreaves, E., Hawkins, R.D., Kandel, E.R., Shapiro, M., and Muller, R.V. (1998). Abolition of long-term stability of new hippocampal place cell maps by NMDA receptor blockade. Science 280, 2121–2126. 10.1126/science.280.5372.2121.

57. Muzzio, I.A., Levita, L., Kulkarni, J., Monaco, J., Kentros, C., Stead, M., Abbott, L.F., and Kandel, E.R. (2009). Attention Enhances the Retrieval and Stability of Visuospatial and Olfactory Representations in the Dorsal Hippocampus. PLOS Biology 7, e1000140. 10.1371/journal.pbio.1000140.

58. Dhawale, A.K., Poddar, R., Wolff, S.B., Normand, V.A., Kopelowitz, E., and Ölveczky, B.P. Automated long-term recording and analysis of neural activity in behaving animals. eLife 6, e27702. 10.7554/eLife.27702.

59. Sheintuch, L., Geva, N., Deitch, D., Rubin, A., and Ziv, Y. (2023). Organization of hippocampal CA3 into correlated cell assemblies supports a stable spatial code. Cell Rep 42, 112119. 10.1016/j.celrep.2023.112119.

60. Kesner, R.P., and Hunsaker, M.R. (2010). The temporal attributes of episodic memory. Behav Brain Res 215, 299–309. 10.1016/j.bbr.2009.12.029.

61. Leão, R.N., Mikulovic, S., Leão, K.E., Munguba, H., Gezelius, H., Enjin, A., Patra, K., Eriksson, A., Loew, L.M., Tort, A.B.L., et al. (2012). OLM interneurons differentially modulate CA3 and entorhinal inputs to hippocampal CA1 neurons. Nat Neurosci 15, 1524–1530. 10.1038/nn.3235.

62. Hasselmo, M.E., and McGaughy, J. (2004). High acetylcholine levels set circuit dynamics for attention and encoding and low acetylcholine levels set dynamics for consolidation. In Progress in Brain Research Acetylcholine in the Cerebral Cortex. (Elsevier), pp. 207–231. 10.1016/S0079-6123(03)45015-2.

63. Haam, J., Zhou, J., Cui, G., and Yakel, J.L. (2018). Septal cholinergic neurons gate hippocampal output to entorhinal cortex via oriens lacunosum moleculare interneurons. Proc. Natl. Acad. Sci. U.S.A. 115. 10.1073/pnas.1712538115.

64. Bell, L.A., Bell, K.A., and McQuiston, A.R. (2015). Activation of muscarinic receptors by ACh release in hippocampal CA1 depolarizes VIP but has varying effects on parvalbumin-expressing basket cells. J Physiol 593, 197–215. 10.1113/jphysiol.2014.277814.

65. Stefanelli, T., Bertollini, C., Lüscher, C., Muller, D., and Mendez, P. (2016). Hippocampal Somatostatin Interneurons Control the Size of Neuronal Memory Ensembles. Neuron 89, 1074–1085. 10.1016/j.neuron.2016.01.024.

66. Sheffield, M.E., and Dombeck, D.A. (2019). Dendritic mechanisms of hippocampal place field formation. Current Opinion in Neurobiology 54, 1–11. 10.1016/j.conb.2018.07.004.

67. Aronov, D., and Tank, D.W. (2014). Engagement of Neural Circuits Underlying 2D Spatial Navigation in a Rodent Virtual Reality System. Neuron 84, 442–456. 10.1016/j.neuron.2014.08.042.

68. Urai, A.E., Aguillon-Rodriguez, V., Laranjeira, I.C., Cazettes, F., The International Brain Laboratory, Mainen, Z.F., and Churchland, A.K. (2021). Citric Acid Water as an Alternative to Water Restriction for High-Yield Mouse Behavior. eNeuro 8, ENEURO.0230-20.2020. 10.1523/ENEURO.0230-20.2020.

69. Cai, C., Friedrich, J., Singh, A., Eybposh, M.H., Pnevmatikakis, E.A., Podgorski, K., and Giovannucci, A. (2021). VolPy: Automated and scalable analysis pipelines for voltage imaging datasets. PLOS Computational Biology 17, e1008806. 10.1371/journal.pcbi.1008806.

70. Pnevmatikakis, E.A., and Giovannucci, A. (2017). NoRMCorre: An online algorithm for piecewise rigid motion correction of calcium imaging data. Journal of Neuroscience Methods 291, 83–94. 10.1016/j.jneumeth.2017.07.031.

71. Grijseels, D.M., Shaw, K., Barry, C., and Hall, C.N. (2021). Choice of method of place cell classification determines the population of cells identified. PLoS Comput Biol 17, e1008835. 10.1371/journal.pcbi.1008835.

72. Skaggs, W., McNaughton, B., and Gothard, K. (1992). An Information-Theoretic Approach to Deciphering the Hippocampal Code. In Advances in Neural Information Processing Systems (Morgan-Kaufmann).

